# The lateral entorhinal cortex is a hub for local and global dysfunction in pre-tauopathy states

**DOI:** 10.1101/2020.04.03.022590

**Authors:** Francesca Mandino, Ling Yun Yeow, Renzhe Bi, Lee Sejin, Han Gyu Bae, Seung Hyun Baek, Chun-Yao Lee, Hasan Mohammad, Chai Lean Teoh, Corey Horien, Jasinda H. Lee, Mitchell K. P. Lai, Sangyong Jung, Yu Fu, Malini Olivo, John Gigg, Joanes Grandjean

**Author notes:** **Corresponding author** Joanes Grandjean, PhD, Department of Radiology and Nuclear Medicine & Donders Institute for Brain, Cognition, and Behaviour, Donders Institute, Radboud University Medical Centre, Nijmegen 6525 EZ, The Netherlands.

## Abstract

Functional network activity alterations are one of the earliest hallmarks of Alzheimer’s disease (AD), detected prior to amyloidosis and tauopathy. Better understanding the neuronal underpinnings of such network alterations could offer mechanistic insight into AD progression. Here, we examined a mouse model (early-tauopathy 3xTgAD mice) recapitulating this early AD stage. We found resting functional connectivity loss within ventral networks, including the entorhinal cortex, aligning with the spatial distribution of tauopathy reported in humans. Unexpectedly, in contrast to decreased connectivity at rest, 3xTgAD mice show enhanced fMRI signal within several projection areas following optogenetic activation of the entorhinal cortex. We corroborate this finding by demonstrating neuronal facilitation within ventral networks and synaptic hyperexcitability in projection targets. 3xTgAD mice thus reveal a dichotomic hypo-connected resting/hyper-responsive active phenotype. The strong homotopy between the areas affected supports the translatability of this pathophysiological model to tau-related deficits in humans.

## Introduction

Tauopathies are a hallmark of many neurodegenerative disorders, including Alzheimer’s disease (AD) ^1^. To date, clinical trials targeting later stages of AD pathology have all failed (except the recently approved aducanumab ^2^), or, in some cases, made symptoms worse ^3^. As such, there is increased interest in identifying predictive biomarkers at the earliest stages of illness, before symptoms have become clinically apparent. Synaptic dysfunction is one such candidate ^4^, and is thought to lead to a discrepancy in resting vs. evoked functional activity patterns in early AD. For example, decreased connectivity patterns are typically observed at rest, while increased connectivity is observed during tasks in early AD patients (^5–8^).

Despite a strong interest in investigating the early stages of illness, the mechanisms underlying aberrant connectivity reorganization are not known. Transgenic animals bearing mutations from familial AD and tauopathies also develop several of the distributed network dysfunctions found in AD, such as in the early stages of cerebral amyloidosis ^9–11^. To understand the physiological basis underlying discrepant resting vs. evoked brain activity patterns during pre-tauopathy, we used the triple-transgenic mouse model for AD (3xTgAD) ^12^. 3xTgAD mice were positive for phospho-tau, a precursor of neurofibrillary tangles, a central feature of tauopathy. Specifically, phospho-tau accumulation was found in the amygdala by 3 months of age, and in the hippocampus at 6 months of age. We further found that local connectivity loss at rest resulted in macroscale network dysfunction already by the 3 month time point. The spatial distribution of these deficits showed a high degree of overlap with homologous networks affected by tauopathies in humans ^13^.

Finally, we employed optogenetics in combination with fMRI (ofMRI) to examine the evoked response from photostimulation of the lateral entorhinal cortex (ENTl), a central dysfunctional node in 3xTgAD ^14,15^ and AD patients ^16–18^. Unexpectedly, this revealed a hyperemic response in 3xTgAD relative to wild-type mice in several distal projection areas.

Our observations underscore several of the physiological underpinnings behind local and distal connectivity dysfunctions commonly observed in groups at risk of developing AD and other tauopathies, thus supporting a reconciliation for the apparent discordant results put forward in early-AD subjects ^5,19,20^. This study provides a neurophysiological model of early network disturbances in AD and points to key translational targets of clinical interest.

## Results

### Ventral networks are affected during tauopathy

To identify the networks affected in tauopathy, we first performed a neuroimaging literature meta-analysis for the search term ‘tauopathy’ through the NeuroQuery library, based on loci reported in 70 studies ^21^. A convergence of loci was found in ventral networks, including the entorhinal cortex, the amygdala, and the nucleus accumbens (Figure 1a), corresponding with the areas affected in the early Braak stages of the pathology ^13,22^.

**Figure 1.**
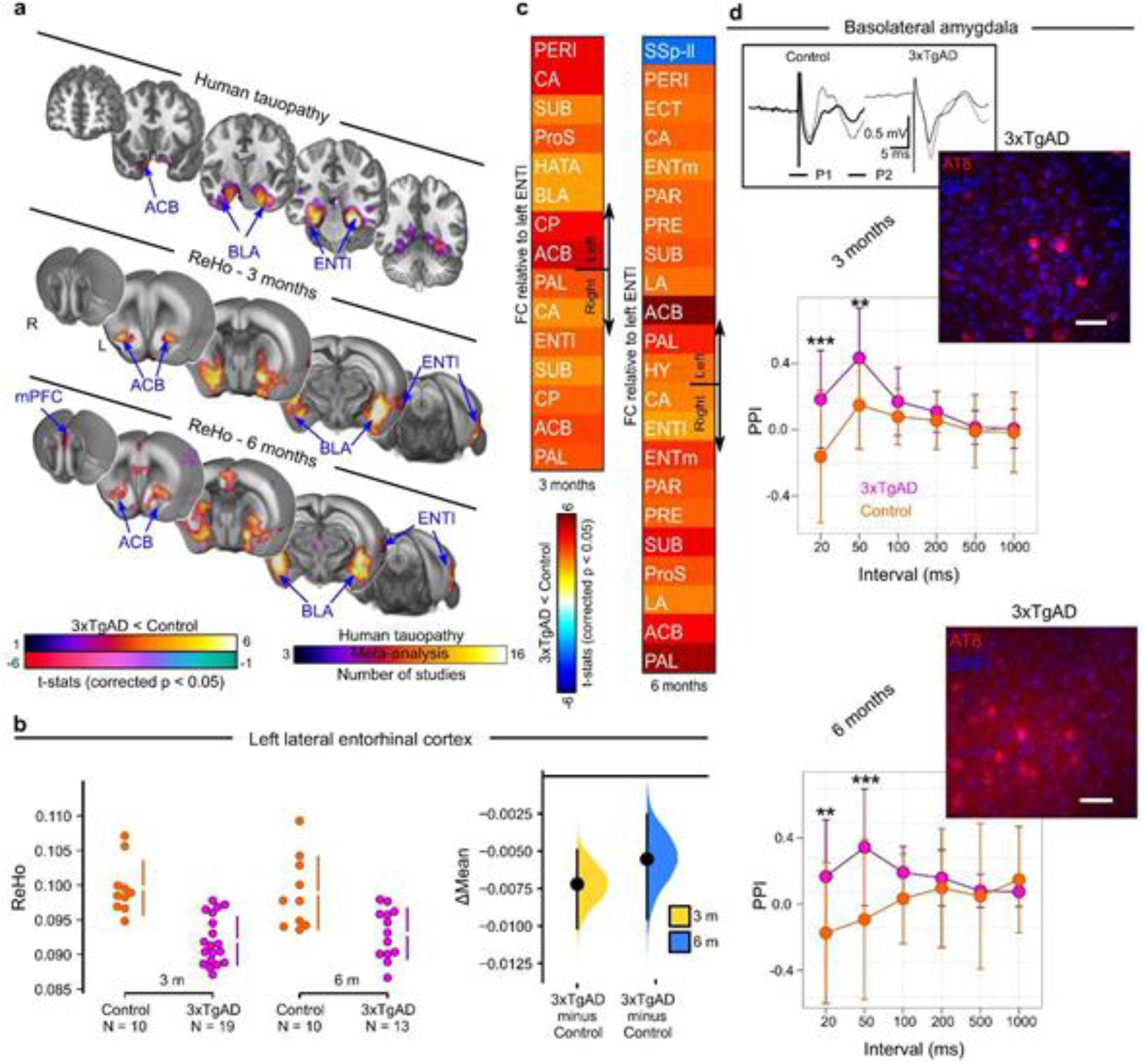
Loss of functional connectivity at rest, and enhanced evoked response, in 3xTgAD. **a)** Networks implicated in human tauopathy as revealed by meta-analysis (top) and Regional Homogeneity (ReHo) thresholded group analysis for 3- and 6-month-old mice (middle, bottom). Significant decrease in ReHo within the ENTl, ACB, and BLA in 3- and 6-month-old 3xTgAD, respectively. **b)** ReHo distribution in the ENTl (left hemisphere: Δmean_3months_ = -0.007 [-0.010, -0.005]; Δmean_6months_ = -0.006 [-0.009, -0.003]). **c)** Pair-wise ROI interactions relative to the left ENTl: decrease of functional connectivity in contralateral ENTl, ACB, BLA at both ages. Increased functional connectivity in somatosensory regions. **d)** Paired-pulse electrical stimulation of the ENTl is recorded as fEPSPs in the BLA: increased facilitation in 3xTgAD, at both ages. Response to ENTl stimulus pairs calculated as the paired-pulse index (PPI) was reported for intervals of 20, 50, 100, 200, 500 and 1000 ms. * p < 0.05, ** p < 0.005, ***p < 0.001 (3 months old: N_controls_ = 7, N_3xTgAD_ = 6; 6 months old: N_controls_ = 4, N_3xTgAD_ = 4). Immunohistochemistry (AT8/DAPI) reveals phosphorylated tau in the BLA, at both ages (age-matched insets). Top-left inset: example of raw data for control and 3xTgAD in BLA for pulse 1 (P1) and pulse 2 (P2) at 3 months. SSp-II: somatosensory area, lower limb; ENTl: lateral entorhinal cortex; BLA: basolateral amygdala; ACB: nucleus accumbens; fEPSP: field excitatory postsynaptic potential. Scale bar: 100 μm

To understand the physiological processes underlying network dysfunction during tauopathy, we turned to the 3xTgAD model of cerebral amyloidosis and tauopathy. To examine early pathological processes, we studied mice at a stage corresponding to Braak ∼II in humans, namely phospho-tau in entorhinal-hippocampal-ventral areas ^13^. Brain slices from 3xTgAD and control animals aged 3, 6, and 10 months were examined for immunoreactivity to 6e10 (targeting the N-terminus of beta-amyloid, Aβ) and AT8 (targeting neurofibrillary tangle-specific phospho-tau (Ser202, Thr205), see Additional file 1: Supplementary Method).

Consistent with previous work ^12,23^, no immunoreactivity was observed to 6e10 at 3 and 6 months of age (Additional file 1: Figure S1), while AT8 binding was revealed in the amygdala at 3 months and also observed in the hippocampus at 6 and 10 months (Figure 1d, Additional file 1: Figure S2). Only at 10 months of age did 3xTgAD exhibit the immunoreactivity patterns attributable to neurofibrillary tangles (Additional file 1: Figure S2c).

We confirmed these results with ELISA performed on brain tissue homogenate. Soluble amyloid peptides levels in 3xTgAD were of comparable magnitude (∼12000/∼16000 pg/mg) to that of age-matched control animals at 3 months of age; a difference of 1.7-fold in soluble amyloid peptides was seen between controls (∼9000 pg/ml) and 3xTgAD (∼16000 pg/ml) at 6 months of age. No difference was observed for insoluble amyloid peptides at both age points. Furthermore, 3xTgAD mice showed an increase in tau levels of several orders of magnitude compared to age-matched controls at both age points, e.g., 20-fold increment at 6 months of age. We conclude that 3xTgAD mice aged 3 and 6 months represent a pre-plaque and pre-tangle stage of AD-like pathology, equivalent to Braak stage ∼II. Therefore, the 3- and 6-months age points were examined for the remainder of this study.

### Functional deficits in 3xTgAD ventral network during pre-tauopathy stage

To examine spontaneous fluctuations in brain activity, we recorded rsfMRI in male 3xTgAD and wild-type control mice on the same background strain (129sv/c57bl6) at 3 (N_3xTgAD_ = 19, N_controls_ = 10) and 6 months of age (N_3xTgAD_ = 13, N_controls_ = 10), longitudinally. The rsfMRI protocol employed here was recently compared to others in a multicenter study, which indicated elevated sensitivity and specificity for resting-state networks detected in this dataset relative to other protocols, including an awake mouse protocol ^24^. One 3xTgAD mouse developed hydrocephalus, which, despite the acute condition, only marginally affected functional connectivity ^25^. This animal was removed from our study following *a priori* exclusion criteria.

Firstly, we examined local connectivity coherence using the Regional Homogeneity (ReHo) method ^26^, a sensitive indicator of local connectivity in mice ^27^. 3xTgAD mice in both age groups presented a bilateral deficit in ReHo, compared to controls, localized to the ventral-amygdaloid-striatal system (Figure 1a). The latter included the ENTl (Δmean_3months_ = -0.007 [-0.010, -0.005]; Δmean_6months_ = -0.006 [-0.009, -0.003], Figure 1b) within the retrohippocampal area, the nucleus accumbens (ACB; Additional file 1: Figure S3a) within the ventral striatum, basolateral amygdala (BLA; Additional file 1: Figure S3b) and medial prefrontal cortex (mPFC, prelimbic cortex within the mPFC reported as an example in Additional file 1: Figure S3c). Concurrently, 3xTgAD mice exhibited increased ReHo in somatosensory areas, such as barrel field cortex (SSp-bfd, Δmean_3months_ = 0.122 [0.005, 0.020], Δmean_6months_ = 0.017 [0.006, 0.035], Additional file 1: Figure S3d).

Strikingly, the functional deficit hotspots, identified with the ReHo method, overlapped with homologous areas in the human brain, namely, the ventral-amygdaloid-striatal system. These areas represent an early target for tau aggregation, as consistently highlighted in a meta-analysis for the keyword ‘tauopathy’ through the NeuroQuery library (Figure 1a). Thus, the 3xTgAD model, similar to other transgenic models, e.g. PSAPP, ArcAβ ^9,28,29^, presents functional alterations that precede extracellular Aβ and tangle deposition and aligns with the spatial distribution of pathology reported in patients affected by mild cognitive impairment. Importantly, deficits within the ventral-amygdaloid-striatal system are consistent with behavioral results in young 3xTgAD, where fear and emotional processes are highly affected ^30^. Emotional control, regulated by the hippocampal-prefrontal-amygdaloid system, is also affected in pre-AD patients ^30,31^, further highlighting the trans-species relevance of our results. Moreover, not all brain areas were affected in a comparable manner: the somatosensory cortex of 3xTgAD presented elevated ReHo, reminiscent of previous findings in the APP transgenic model ^32^. This highlights that pathophysiology does not affect each brain region equally.

### Dopamine response genes are enriched in functionally compromised regions in 3xTgAD

We hypothesized that the patterns of functional alteration were the consequence of 3xTgAD transgene products interacting with others from the brain transcriptome. To test this, we searched for gene expression patterns overlapping with the functional deficits highlighted above (Figure 1a). The expression profile from 4117 genes selected for their brain expression was spatially correlated with the ReHo deficits in the 3-month-old 3xTgAD dataset (Additional file 1: Figure S4). We compared the occurrence in biological processes in a ranked-test in the GOrilla database. Genes associated with dopamine signaling overlapped with regional deficits (Table 1, p-value_FDR_ < 0.001). These included genes encoding for G protein signaling (Rgs9, EntrezID = 19739, r = 0.178), G protein subunit (Gnal, EntrezID = 14680, r = 0.215), and Adenylate cyclases (Adcy5, EntrezID=224129, r = 0.184, Adcy6, EntrezID = 11512, r = 0.142).

**Table 1.**
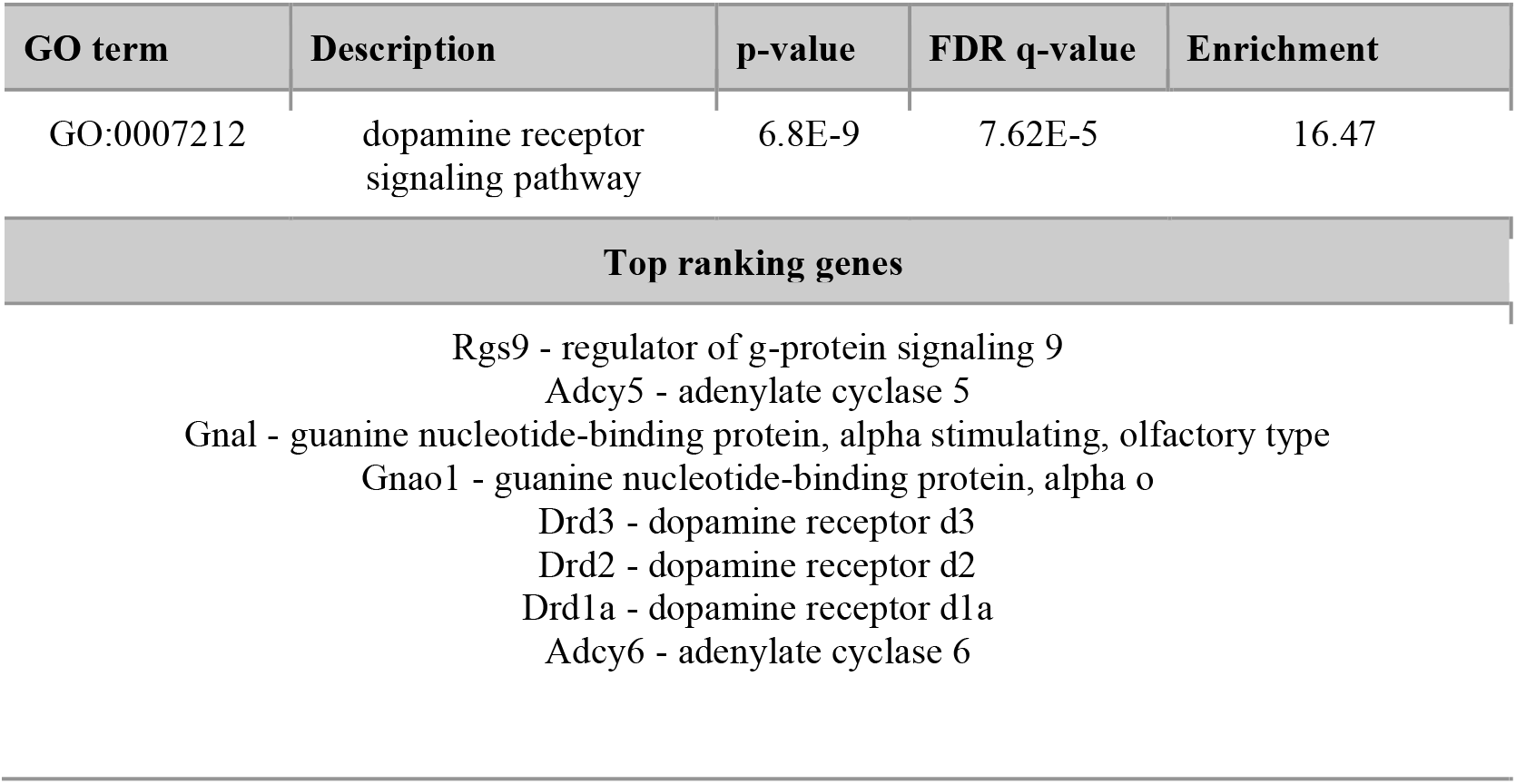
Dopamine receptor signaling pathway genes overlap with the ReHo functional deficit in 3xTgAD.

Changes in dopamine signaling were reported previously in an animal model of cerebral amyloidosis also overexpressing APP_swe_. In addition, loss of midbrain dopamine (DA) neurons, as well as deficits in hippocampus-to-ACB signaling mediated by DA, has been observed at 3 months of age in Tg2576 mice ^33,34^. In AD, alterations in DA levels or DAergic receptors are found to significantly impact synaptic plasticity and hippocampal-memory encoding ^35^. Loss of DA receptors, especially D2, has been shown in areas such as the hippocampus, prefrontal cortices, and BLA ^36,37^ in AD patients. PET studies on AD patients also confirm a loss of striatal D2-like receptors ^38^. Our results, therefore, bring supporting evidence for an interaction between the DAergic system associated with early cerebral amyloidosis and tauopathy, which leads to synaptic dysfunction.

### Local functional connectivity deficits translate into whole-brain network alterations

The ENTl and associated hippocampal areas are fundamental for declarative memory encoding and retrieval ^39,40^. In particular, the ENTl is among the first hubs affected in both human AD ^41^ and the 3xTgAD model of cerebral amyloidosis and tauopathy ^12^ (Figure 1a). As such, the ENTl was targeted for further analysis. To examine distal functional connectivity alterations at rest, we assessed pair-wise ROI interactions relative to the left-hemisphere ENTl. Functional connectivity to the ENTl was decreased in the retrohippocampal and hippocampal regions (Figure 1c, and e.g. ENTl right hemisphere: Δmean_3months_ = -0.114 [-0.193, -0.035], Δmean_6months_ = -0.127 [-0.205, -0.058], Additional file 1: Figure S5c), ventral striatum (ACB, Additional file 1: Figure S5b), and amygdala (BLA, Additional file 1: Figure S5c). Similarly, an increase in functional connectivity relative to the ENTl was reported in the somatosensory cortex (lower limb, SSp-ll; Additional file 1: Figure S5d). These results, focusing on the ENTl-specific network, show similarities to the whole-brain functional connectivity changes assessed with ReHo, presented in an overlapped design in Additional file 1: Figure S6.

To confirm the connectivity results, we performed electrophysiological recordings in urethane-anesthetized 3xTgAD and control mice, *in vivo*, at 3 or 6 months of age (Additional file 1: Supplementary Method). Field excitatory postsynaptic potentials (fEPSP) were assessed within the BLA and dentate gyrus (DG), following electrical stimulation in the ENTl. A paired-pulse stimulation (PPS) protocol, used to assess short-term synaptic plasticity changes, was analyzed through the paired-pulse index (PPI) and revealed a quadruple effect between ROI, age, genotype, and paired-pulse intervals: F_(51,46)_ = 28.135, p < 0.001. Specifically, no difference between 3xTgAD and controls was found for longer paired-pulse intervals in both the BLA and DG at both age points. A strong increase in facilitation was instead reported for short intervals in both BLA (e.g. 3 months old: 20 ms, z-score = -3.96, p < 0.001; 50 ms, z-score = -3.11, p < 0.05; Figure 1d) and DG (e.g. 3 months old: 20 ms, z-score = -3.16, p < 0.01; 50 ms, z-score = -4.54, p < 0.001; Additional file 1: Figure S7). This may indicate neuronal synaptic hyperexcitability in 3xTgAD: intracellular Ca^2+^ residuals from the first stimulus (P1) likely elicit augmented release of the presynaptic neurotransmitter in response to the second pulse (P2). This hyperexcitable neuronal profile may support the network dysfunction observed, through compensatory mechanisms in response to the compromised functional connectivity reported at rest. Additionally, the Input/Output curve (IOC) analysis for synaptic strength revealed a quadruple interaction effect between ROIs, age, genotype, and stimulation amplitude: F _(35,32)_ = 108.31, p < 0.001. Within the BLA, 3xTgAD mice did not show significant differences compared to controls at 3 months, whereas, there was a significantly larger response at 6 months for all stimulus intensities, e.g. 300 μA (z = -6.23, p < 0.001), 450 μA (z = -6.64, p < 0.001) and 600 μA (z = -7.1, p < 0.001; Additional file 1: Figure S8a). Within DG, 3xTgAD mice showed larger responses than controls by 3 months (e.g. 300 μA z-score = 2.36, p < 0.05 and 600 μA z-score = 2.93, p < 0.005; Additional file 1: Figure S8b, left panel). At 6 months, there was a significant difference between genotypes for the strongest current stimulus (600 μA, z-score = 2.12, p < 0.05), although there was a trend for increased facilitation in 3xTgAD mice compared to controls (Additional file 1: Figure S8b, right panel). Taken together, our electrophysiological *in-vivo* evidence reveals hyperexcitable behavior during the evoked neuronal response in disease-relevant 3xTgAD brain regions. This suggests a dichotomous relationship between increased-evoked and reduced-spontaneous activity in AD-like vulnerable areas, where functional connectivity is highly compromised at rest.

In an exploratory analysis, we examined whole-brain network deficits in 3xTgAD mice at 3 and 6 months of age (Additional file 1: Figure S9b). Alterations were consistent between both age groups and localized within and between regions highlighted in the ReHo analysis, namely, in the amygdaloid/cerebral nuclei (including BLA and ACB) and the ENTl and the hippocampal formation (Figure 1a, Additional file 1: Figure S9b). Importantly, the nodal degree distribution, i.e., the number of affected connections per ROI, was found to overlap with ReHo (Additional file 1: Figure S9b). This indicates that local functional connectivity deficits translate into distal functional connectivity deficits and, in turn, greater network dysfunction. Moreover, regions highlighted in the pairwise correlation analysis (amygdala and hippocampus, Additional file 1: Figure S2) were also found to be enriched in tau aggregates consistent with tau dispersions across functionally connected networks ^42^.

### Functional connectome of the ENTl revealed by optogenetics

The ENTl was revealed above to be a major hub region affected in the 3xTgAD brain at rest. To further explore the functional consequences of this finding,, we leveraged fMRI combined with optogenetics (ofMRI; Figure 2) ^43^ to visualize the hemodynamic response to a 10-block design photostimulation of ChR2-transfected CaMKIIα-positive (AAV5-CamKIIα-hChR2 (H134R)-mCherry) ENTl neurons (N_controls_ = 10, N =12_3xTgAD_; Figure 2ab, Additional file 1: Figure S10ab). Anatomical imaging of the optic fiber revealed that the ENTl was accurately targeted (Figure 2c, Additional file 1: Figure S10c). Transfection led to robust expression at the target site (Figure 2b, Additional file 1: Figure S11a). Cell bodies of transfected neurons were consistent with those of excitatory pyramidal cells (Additional file 1: Figure S11b, Supplementary Method). ENTl neurons faithfully responded to 5 and 20 Hz light pulse trains in both controls and 3xTgAD (Figure 2d, Additional file 1: Figure S11c). Prolonged photostimulation (500 ms) applied to patched neurons *in vitro* confirmed an effective ChR2-induced inward current on both control and 3xTgAD mice *post hoc* (Additional file 1: Figure S11d).

**Fig.2.**
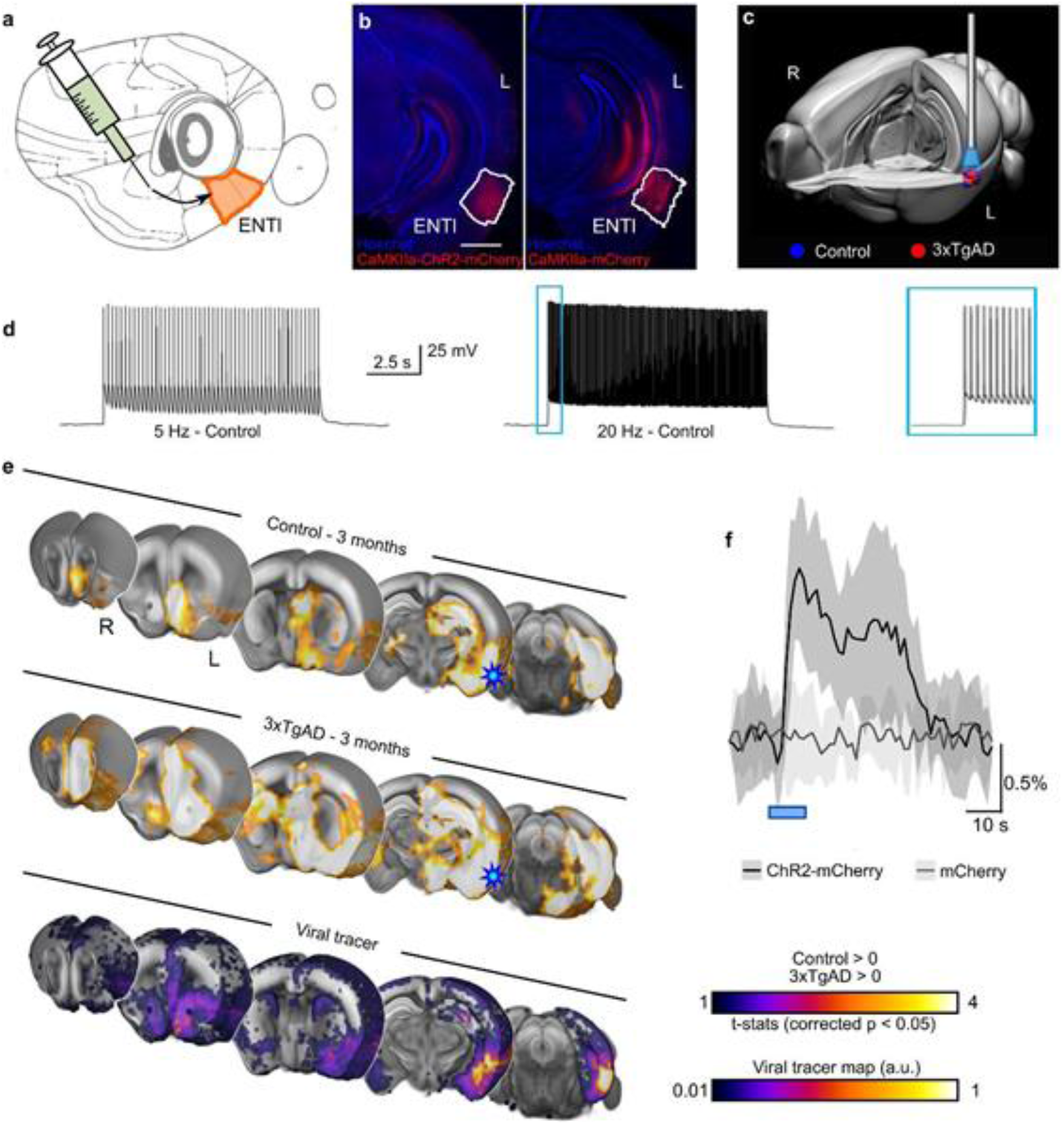
Optogenetic modulation of the lateral entorhinal cortex. **a)** Diagram of stereotaxic injection in the left ENTl (in mm from Bregma and middle line: AP: -2.8, ML: +4.2, DV: -2.6). **b)** ChR2-mCherry/Hoechst (left) and mCherry/Hoechst (right) indicate successful targeting of the ENTl and stable transfection. **c)** 3-d rendering of fiber tip position for each experimental animal, red dots indicate 3xTgAD subjects (N = 12) and blue dots indicate controls (N = 10). **d)** Optogenetic stimulation of ChR2-transfected neurons at 5 Hz and 20 Hz *in vitro* shows frequency-locked spiking activity. **e)** Optogenetically-locked BOLD response overlaps with density projection maps. Top and middle sections indicating one-sample t-test for stimulation-locked BOLD response in the control group and 3xTgAD mice (N = 10, N = 12, respectively; p < 0.05 corrected) highlighting activation in key regions related to ENTl projections, i.e., HIP, BLA, ACB and mPFC; AIBS tracer projection density map with injection in ENTl (experiment ID: #114472145; bottom section). Stimulation site (ENTl) indicated with a blue star. **f)** BOLD response profile, averaged across 10 blocks, following 20 Hz stimulation in control mice injected with ChR2-mCherry (N = 10) and control mice injected with mCherry only (N = 9) shows an opsin-dependent BOLD response. AP: anterior-posterior; ML: medio-lateral; DV: dorso-ventral; HIP: hippocampus; BLA: basolateral amygdala; ACB: nucleus accumbens; mPFC: medial prefrontal cortex. Scale bar: 1000 μm

There was no evidence of aberrant spontaneous behavior to photostimulation protocols in freely behaving mice, unlike seizures previously reported following photostimulation of the hippocampus in rats ^44^. An unbiased voxel-wise analysis revealed the BOLD signal associated with our modeled response in controls and 3xTgAD at 3 months of age (Figure 2e, top and middle rows respectively) in several regions, including the hippocampal formation (hippocampus and retro hippocampal areas), the amygdaloid area (e.g. BLA), the ventral striatum (ACB), the prefrontal and the insular areas. Similar results were reported for the 6-month age point (Additional file 1: Figure S12, left and middle panels). Interestingly, optogenetically-evoked activity was mostly confined to the ipsilateral hemisphere, despite the presence of contralateral projections, as predicted by viral tracers (Figure 2e bottom row, spatial correlation r = 0.36). This supports the notion of a neuronal mechanism that silences the response contralaterally but not ipsilaterally to artificially generated neuronal activity, perhaps via feed-forward active inhibition ^45^. The response elicited through photostimulation of ENTl was comparable between mice at 3 and 6 months of age, indicating a stable expression allowing for longitudinal analysis. A negative control carried out in healthy wild-type mice (N_mCherry-controls_ = 9) transfected with mCherry alone (Figure 2b) did not reveal the presence of a light response, except for a visual-associated response of the lateral geniculate nucleus and superior colliculus, probably due to the fiber illumination received as a direct visual response to retinal illumination (Additional file 1: Figure S10d). Hence, we concluded that the response recorded with ofMRI was not associated with potential heating and/or vascular photoreactivity artifacts. Photostimulation at frequencies ranging from 5 to 20 Hz indicated spatially overlapping results (Additional file 1: Figure S10e), in contrast to previous research ^46^. In fact, the non-specific, visually-associated response amplitude was stronger at lower frequencies (Additional file 1: Figure S10d, upper panel), while the opsin-associated response was more marked at 20 Hz (Figure 2e, Additional file 1: Figure S12). The areas associated with a visual response elicited with 5 Hz stimulation were subsequently masked from our results and the remainder of the analysis focused on the 20 Hz paradigm (Additional file 1: Figure S10d, lower panel).

### Potentiated hemodynamic response and neuronal activity in 3xTgAD

To assess response differences across the brain, a non-parametric second-level analysis comparing the amplitude of activation between 3xTgAD and controls was carried out (Figure 3, central segment). Group size at 6 months of age was reduced due to group attrition, e.g., detachment of the implant. Therefore, group sizes differed between 3 months (N_controls_ = 10, N_3xTgAD_ = 12) and 6 months (N_controls_ = 8, N_3xTgAD_ = 10) of age. Surprisingly, and in contrast to the results observed at rest, 3xTgAD mice showed significantly larger responses across several regions compared to controls at 3 months. The regions affected included the ipsilateral dorsal hippocampus, ACB, mPFC, cingulate and retrosplenial areas, and contralateral ENTl. The presence of an effect at 6 months could not be detected, potentially due to the reduced group size, or to a normalization of the response at a later age.

**Figure 3.**
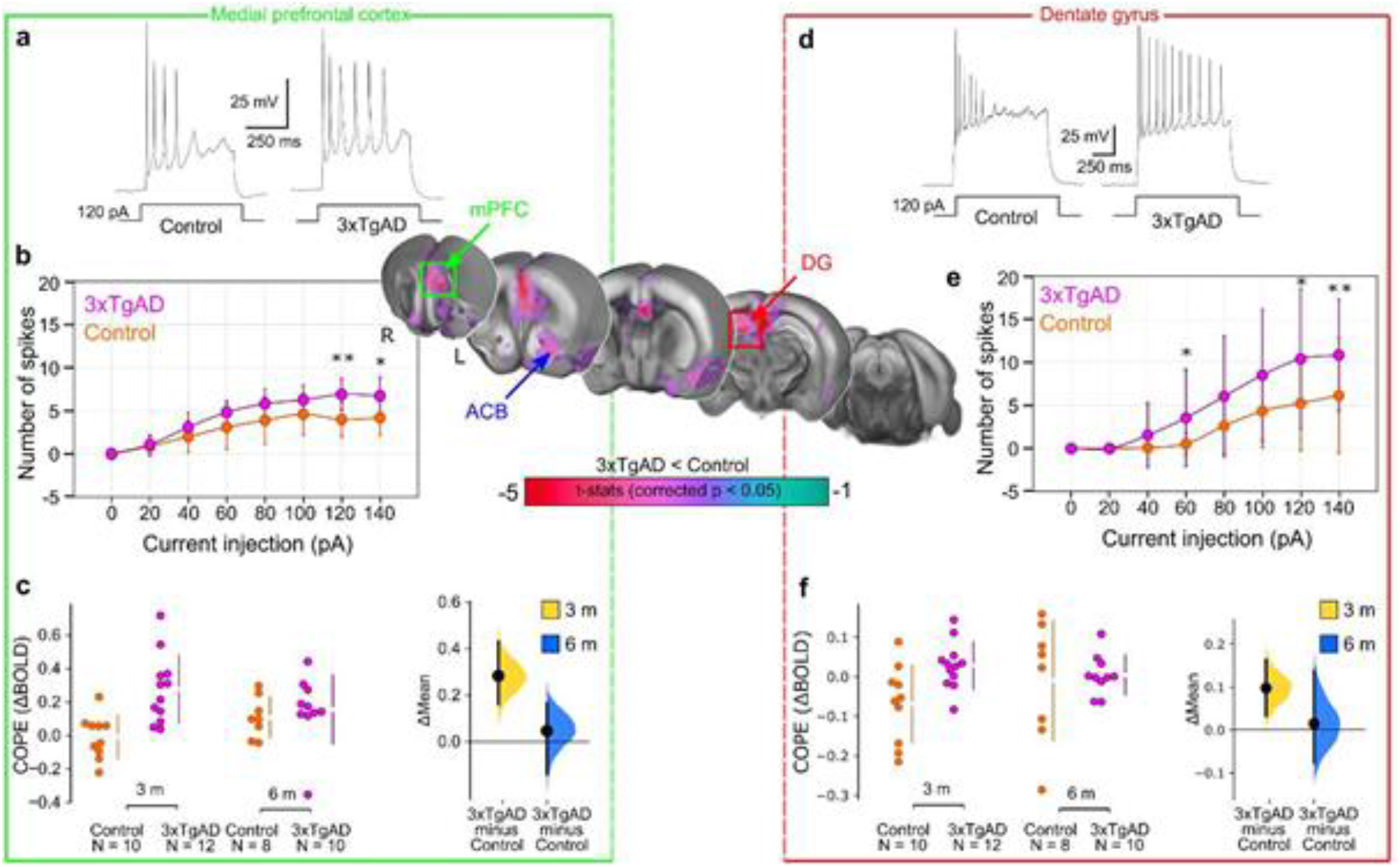
Increased optogenetically-locked response in 3xTgAD compared to controls at 3 months. Central panel: Two-sample t-test showing significantly higher response (p < 0.05, corrected) in 3xTgAD compared to controls in AD-like vulnerable regions, such as mPFC and DG, left and right sections respectively. **a**,**d)** Representative action potential firing patterns to 120 pA injection in ILA pyramidal cells (N_controls_ = 4, n = 10 / N _3xTgAD_ = 4, n = 11) and DG granule cells (N_controls_ = 4, n = 29 / N _3xTgAD_ = 5, n = 18) respectively. **b, e)** The evoked spike number during various injected currents in ILA pyramidal cells and DG granule cells, respectively. The data are plotted as mean action potential numbers ± 1 SD. The statistical significance is presented with asterisks (*p < 0.05, **p < 0.01 by Mann-Whitney U test). **c, f)** COPEs are represented for 3xTgAD and controls at both age points for mPFC and DG, respectively

To examine local response amplitude, contrast of parameter estimates (COPEs) were extracted from ROIs highlighted in the voxel-wise comparison: ENTl (Δmean_3months_ = 0.044, [0.022, 0.070]), ACB (Δmean_3months_ = 0.15 [0.0106, 0.298]), mPFC (Figure 3c, Δmean_3months_ = 0.283 [0.161, 0.431]) and DG (Figure 3f). Similar results were present at lower stimulation frequencies, although the more distal regions had a mitigated effect at lower frequencies (example for 10 Hz, ENTl: Δmean_3months_ = 0.008, [-0.08, 0.09]). Data acquired at rest, prior to optogenetic stimulation, indicated ReHo deficits converging with that acquired in the previous dataset, replicating our observations above, despite lower acquisition quality due to a room temperature receiver coil instead of a cryoprobe coil (Additional file 1: Figure S13).

To confirm the increased response observed in ofMRI, we examined neuronal excitability *ex vivo* in two projection areas highlighted above; the DG, within the hippocampus, and the infralimbic area (ILA), within mPFC, (Additional File 1: Supplementary Method). Acute brain slice electrophysiology indicated that excitatory neurons of 3xTgAD, in both ILA (Figure 3ab) and DG (Figure 3de), were prone to increase the number of spikes derived by current injection compared to controls, but ILA neurons showed enhancement of afterhyperpolarization (AHP) latency and half-width of the action potential, a phenotype consistent with previous results ^47^ (Additional file 1: Table S2), while spike morphology was comparatively unaltered (Additional file 1: Table S3). To examine how the alterations of the ENTl functional projectome at 3 months related to a loss of functional connectivity at rest, we projected the two responses onto the same template (Additional file 1: Figure S12, right panel). We found that several of the ROIs, presenting decreased functional connectivity at rest, were responding with greater amplitude to optogenetically-driven neuronal activity. This was the case for regions encompassing the ventral striatum (ACB), the dorsal DG within the hippocampal formation and prefrontal regions.

In sum, 3xTgAD mice showed decreased functional connectivity at rest and, in contrast, an increased response during optogenetic stimulation. Direct neuronal recordings, both in vivo and ex vivo, showed a hyperactive phenotype. Taken together, these results support a hypo-connected/hyper-responsive dichotomy characterizing rest vs. evoked states in early stages of pathology progression in 3xTgAD.

## Discussion

Neurological disorders, including tauopathies such as AD, are one of the greatest challenges in modern medicine. The absence of disease-modifying treatment for AD represents a major loss for the millions of patients affected worldwide. A detailed understanding of the disease mechanisms across spatial and temporal scales constitutes a translational opportunity to facilitate the drug development process. Here, we have determined that the pre-tangle stage of the 3xTgAD mouse model presents functional connectivity patterns overlapping with areas affected by tauopathy in humans. This supports the trans-species relevance of our results.

We determined that distal connectivity disturbances could be explained by local connectivity deficits. Moreover, we observed a dichotomy between resting activity, which leads to decreased connectivity, and evoked activity, that promotes increased metabolic demand and neuronal hyperexcitability. We confirmed fMRI results with electrophysiological recordings that indicated a pathological effect on signal transmission and electrical properties of neurons in vulnerable neuronal circuits. Hence, we demonstrated that fMRI connectivity deficits are rooted in a deeper physiological context. Further, an in-silico transcriptome analysis indicated a potential association of these results with deficits in the DA system. Importantly, our results connect to two widely examined models of AD pathological progression, namely the tau seeding hypothesis ^13,48^, and a revised amyloid hypothesis ^32^. Finally, we examined potential mechanisms underpinning the electrophysiological signatures. Staging the disease progression has been an important question in AD in order to understand mechanisms and to improve diagnostics ^49^.

We observed phospho-tau spreading from 3 to 10 months in 3xTgAD model mice and found that this was associated with long-range connectivity deficits in young model animals. This underpins the notion of a tight coupling between functional connectivity and tau progression ^42^, supported by molecular work by others ^50^. The seeding hypothesis can be further connected to other models of axonal degeneration and inflammation ^51^ and, thus, it provides a coherent description of the pathophysiology process taking place in AD.

The contribution of different Aβ species (soluble oligomers, fibrils, plaques) has been the subject of ardent discussion in the literature. Here, we demonstrate functional connectivity dysfunction in the absence of amyloid plaques and detectable elevated Aβ levels, similar to previous results in other models ^9,10,32^ and in subjects at risk of developing AD ^5,19^. Our results thus contribute to the view that the traditional amyloid cascade hypothesis for AD ^52^ etiopathogenesis should be updated. Additionally, our results contribute to a modern re-interpretation of the cascade and reconcile contradicting results within the human literature. Decreased resting connectivity patterns were paired with an increased response to optogenetically-driven and electrically-driven neuronal activity, thus highlighting a possible increase in neuronal excitability following stimulation and increased metabolic demands (Figure 3, Additional file 1: Figure S12 right panel).

The dichotomy of the direction of these changes in resting and evoked activity, specifically within the ENTl network, mirrors several findings in preclinical and early stages of AD patients. Decreased functional connectivity is found in mild AD patients at rest, in areas related to the DMN, including the hippocampal formation ^53^. However, task-based fMRI studies show increased activity in memory-related areas in subjects at risk of AD but who are cognitively still normal (e.g. APOEε4 carriers) suggesting a dichotomy in network organization of the brain at rest and in engaged status ^54^. Taken together, our findings help to reconcile apparent discordant results put forward in early-AD subjects ^5,7,20^.

Importantly, our results also fit into modern hypotheses for the amyloid cascade. Buckner and colleagues demonstrated that network dysfunction overlapped and preceded amyloid deposition revealed with PET ^55^. Bero and colleagues demonstrated in APP transgenic mouse models that hyper-connectivity patterns at a young age correlated with amyloid plaque distribution later in life ^32^. Our results support the notion that local and distal network dysfunction at rest impairs information transmission and processing. This leads to increased metabolic demand during evoked activity, which putatively leads to circuit exhaustion and further accumulation of Aβ species through increased neuronal activity ^56^. It will be important to confirm this model prediction in older 3xTgAD mice.

## Conclusions

In sum, the functional deficits found within and relative to the temporal and ventral brain areas in 3xTgAD mice recapitulate several important effects described in pre-AD subjects with functional neuroimaging. Importantly, we postulate a disruption in DAergic signaling pathways as one of the earliest features characterizing AD development. Moreover, the photostimulation of the ENTl leads to a marked increase in BOLD response in several important projection areas. The dichotomic behavior between resting and evoked functional responses, taking place during the early stages of cerebral amyloidosis and tauopathy, reveals an endophenotype in line with the human tauopathy profile. This suggests that similar pathophysiological mechanisms might be the cause of network dysfunction in clinical cases, providing an understanding of the underlying mechanisms leading to functional deficits preceding this fatal neurodegenerative disorder.

## Methods

### Animal permit

All procedures conducted in the UK were performed in accordance with the UK Animals (Scientific Procedures) Act 1986 and the University of Manchester Ethical Review Panel under Home Office license PPL 70/7843. All experiments performed in Singapore Bioimaging Consortium, A*STAR, Singapore, were in accordance with the ethical standards of the Institutional Animal Care and Use Committee (A*STAR Biological Resource Centre, Singapore, IACUC #171203).

In both locations, the 3xTgAD and the control colonies were maintained ‘in-house’ through the pairing of homozygous individuals. Mice were housed in cages of up to five, with same-sex and genotype cage-mates in a pathogen-free environment, kept at a 45-65% humidity, under a 12:12-hour light-dark cycle and room temperature, with *ad libitum* access to food and water. The detailed breakdown of animal group sizes per experiment is detailed in Additional file 1: Table S1.

Specifically, male 3xTgAD and control mice on the same background strain (129sv/c57bl6) aged either 3-4 months (N = 6 and N = 7, respectively) or 6-7 months old (N = 4 and N = 4, respectively) were used for electrophysiological recordings *in vivo*. Additionally, male controls (total N = 29) and 3xTgAD (total N = 31) have been used for the imaging experiments. Specifically, N_controls_ = 10 and N_3xTgAD_ = 19 underwent rsfMRI experiments.

Additionally, N_controls_ = 19 and N_3xTgAD_ = 12 mice underwent ofMRI experiments. An *a priori* power analysis was performed following results in ^28^ using R (power.t.test), indicating that N = 10 per group for rsfMRI is sufficient to achieve 80% power with the following parameters: delta = 14, SD = 11, two-tailed test, significance threshold = 0.05.

### Optogenetic surgery

Male 129sv/c57bl6 and 3xTgAD mice (∼30 g, N = 19, N = 12 respectively) were anesthetized with a mixture of ketamine/xylazine (ketamine 75 mg/kg, xylazine 10 mg/kg). The head was shaved and cleaned with three wipes of Betadine® and ethanol (70%). Lidocaine was administered subcutaneously, *in situ* under the scalp. Each animal was kept on a warm pad to prevent hypothermia, and the head was positioned in a stereotaxic frame; protective ophthalmic gel was applied to avoid dryness. A portion of the scalp was removed to expose the skull. The distance between Bregma and Lambda was measured and compared to the standard 4.2 mm reported in the mouse brain atlas. Any deviation from 4.2 mm allowed a proportional adjustment for craniotomy coordinates. Small craniotomies were performed above the left hemisphere with a drill (burr tip 0.9 mm^2^) at -2.8 from bregma, +4.2 from the midline. Virus injection into ENTl was carried out through this craniotomy at -2.8 to -2.7 mm from the brain surface and cannula positioning reached -2.6 mm from the surface.

Coordinates were taken according to the Paxinos mouse brain atlas ^57^. An injection of adeno-associated virus (AAV) was performed in the target location using a precision pump (KD Scientific Inc., Harvard Bioscience) with a 10 μl NanoFil syringe with a 33-gauge beveled needle (NF33BV-2). The AAV used ^58^, AAV5-CaMKIIa-hChR2(H134R)-mCherry (N_controls_ = 10, N_3xTgAD_ = 12) or AAV5-CaMKIIa-mCherry (N_mCherry-controls_ = 9), titer 1-8×10^12^ vg/ml, were acquired from Vector Core at the University of North Carolina (USA). A total volume of 0.75 μl of the vector was injected in each mouse at a rate of 0.15 μl/min. The injector was kept in location for 10 minutes after injection completion to preclude backflow. After the extraction of the needle, a fiber optic cannula (diameter 200 μm, 0.39 NA, length according to the injection site, diameter 1.25 mm ceramic ferrule) was lowered to the targeted region (Laser 21 Pte Ltd, Singapore; Hangzhou Newdoon Technology Co. Ltd, China). The cannula was fixed in place with dental cement (Meliodent rapid repair, Kulzer). Buprenorphine was administered post-surgically to each animal. Animal recovery took place on a warm pad.

### Animal preparation for imaging

Animal preparation followed a previously established protocol ^59^. Anesthesia was induced with 4% isoflurane; subsequently, the animals were endotracheally intubated, placed on an MRI-compatible cradle, and artificially ventilated (90 breaths/minute; Kent Scientific Corporation, Torrington, Connecticut, USA). A bolus with a mixture of Medetomidine (Dormitor, Elanco, Greenfield, Indiana, USA) and Pancuronium Bromide (muscle relaxant, Sigma-Aldrich Pte Ltd, Singapore) was administered subcutaneously (0.05 mg/kg), followed by a maintenance infusion (0.1 mg/kg/hr) administered 5 minutes later while isoflurane was simultaneously reduced and kept to 0.5%. Functional MRI was acquired 20 min following maintenance infusion onset to allow for the animal state to stabilize. Care was taken to maintain the temperature of the animals at 37°C.

### fMRI data acquisition and stimulation protocols

Data were acquired on an 11.75 T (Bruker BioSpin MRI, Ettlingen, Germany) equipped with a BGA-S gradient system, a 72 mm linear volume resonator coil for transmission. A 2×2 phased-array cryogenic surface receiver coil was adopted for the rsfMRI experiment (N = 29) and a 10 mm single-loop surface coil for ofMRI experiments (N = 31). Images were acquired using Paravision 6.0.1 software.

For the rsfMRI data acquisition, an anatomical reference scan was acquired using a spin-echo turboRARE sequence: field of view (FOV) = 17×9 mm^2^, FOV saturation slice masking non-brain regions, number of slices = 28, slice thickness = 0.35, slice gap = 0.05 mm, matrix dimension (MD) = 200×100, repetition time (TR) = 2750 ms, echo time (TE) = 30 ms, RARE factor = 8, number of averages = 2. Functional scans were acquired using a gradient-echo echo-planar imaging (EPI) sequence with the same geometry as the anatomical: MD = 90×60, TR = 1000 ms, TE = 15 ms, flip angle = 50°, volumes = 600, bandwidth = 250 kHz. Parameters for the ofMRI data acquisition were adapted to the lower sensitivity of the room temperature receiver coil. The anatomical reference scan was acquired using FOV = 20×10 mm^2^, number of slices = 34, slice thickness = 0.35, slice gap = 0 mm, MD = 200×100, TR = 2000 ms, TE = 22.5 ms, RARE factor = 8, number of averages = 2. Functional scans were acquired using FOV = 17×9 mm^2^, FOV saturation slice masking non-brain regions, number of slices = 21, slice thickness = 0.45, slice gap = 0.05 mm, MD = 60×30, TR = 1000 ms, TE = 11.7 ms, flip angle = 50°, volumes = 720, bandwidth = 119047 Hz. Field inhomogeneity was corrected using MAPSHIM protocol. Light stimulation was provided through a blue light laser (473 nm, LaserCentury, Shanghai Laser & Optic Century Co., Ltd; ∼12-15 mW output with continuous light at the tip of the fiber) controlled by in-house software (LabVIEW, National Instruments). After an initial 50 s of rest as a baseline, 5, 10 or 20 Hz light pulses (10 ms long pulses) were applied for 10 s followed by a 50 s rest period, in a 10-block design fashion. An additional 60 s of rest were recorded after the last block of stimulation (Additional file 1: Figure S10a). The experimental groups (3xTgAD and wild-type mice with ChR2-mCherry) and the negative control group (wild-type mice with mCherry alone) underwent the same imaging protocol, i.e., one resting-state scan, followed by randomized 5 Hz, 10 Hz and 20 Hz evoked fMRI scans. The negative control group was imaged with the same imaging protocol as the experimental groups to exclude potential heating and/or vascular photoreactivity artifacts ^60,61^. Additionally, in order to exclude abnormal behavior induced by the photostimulation protocol ^44^, all animals underwent the three stimulation sessions (5 Hz, 10 Hz, and 20 Hz) again while awake and freely walking in a behavior-chamber.

### fMRI analysis

Images were processed using a protocol optimized for the mouse and corrected for spikes (*3dDespike, AFNI* ^*62*^), motion (*mcflirt, FSL* ^*63*^) and B1 field inhomogeneity (*fast*). Automatic brain masking was carried out on the EPI using *bet*, following smoothing with a 0.3 mm^2^ kernel (*susan*) and a 0.01 Hz high-pass filter (*fslmaths*). Nuisance regression was performed using FIX ^11^. Separate classifiers were generated for rsfMRI and ofMRI. The EPIs were registered to the Allen Institute for Brain Science (AIBS) reference template ccfv3 using SyN diffeomorphic image registration (*antsIntroduction*.*sh, ANTS* ^*64*^).

Local connectivity was assessed with ReHo (*3dReHo*) ^26,27^. Pair-wise region-of-interest (ROI) analysis was carried out with respect to ROIs defined in the AIBS atlas. Time series extracted with the atlas were cross-correlated to the time series from the ENTl using Pearson’s correlation. The ofMRI response was examined using a general linear model (GLM) framework (*fsl_glm*). The stimulation paradigm and its first derivative were convolved using the default gamma function and used as regressors in the analysis, with motion parameters as covariates. Nomenclature and abbreviations for the brain regions are in accordance with https://atlas.brain-map.org/.

Human literature spatial meta-analysis was performed on the neuroquery.org platform on November, 8^th^ 2019, using the term ‘tauopathy’ in the query (‘https://neuroquery.org/query?text=tauopathy+’). The query returned 30 spatial maps depicting activation voxels in neuroimaging literature associated with the searched term (Figure 1a).

### Anatomical gene expression atlas comparison

The spatial expression profile for 4117 genes was obtained from the anatomical gene expression atlas database using the application programming interface from the AIBS ^65^. The spatial correlation between the ReHo second-level statistical map and each of the genes was estimated using Pearson’s correlation (*fslcc*). ReHo-gene correlations were ranked and tested for enrichment of biological processes using Gene Ontology enRIchment anaLysis and visuaLizAtion tool (GOrilla, http://cbl-gorilla.cs.technion.ac.il/) ^66,67^. Enrichment was tested with Fisher’s Exact test with FDR correction.

### Statistics and data availability

Descriptive statistics for neuroimaging data are given as mean difference and [95th confidence interval] unless stated otherwise, and graphically represented as ‘Gardner–Altman plots’ (https://www.estimationstats.com/; ^68^). If not specified, descriptive statistics are provided for left hemisphere ROIs. The statistical threshold for significance was set at p < 0.05, two-tailed. Voxel-wise was carried out with a non-parametric permutation-based (5000 permutations) test (*randomize*). Cluster correction was carried out with threshold-free cluster enhancement (tfce). Thresholded t-statistic for one-sample and two-sample t-tests (p < 0.05, tfce corrected) are shown as a color-coded overlay on the AIBS template. ROI analysis was carried out with a linear mixed model using genotype and age as fixed effects and individual intercepts as random effects, using the lme4 package (1.1-21) for R (https://cran.r-project.org/, 3.5.3, “Great Truth”). Significance was assessed with general linear hypothesis tests implemented in the multcomp (1.4-10) package and corrected with the false discovery rate.

The rsfMRI and ofMRI datasets supporting the conclusions of this article are available on a CCO license in the OpeNeuro repository, https://openneuro.org/ (DOI: 10.18112/openneuro.ds001890.v1.0.1, 10.18112/openneuro.ds002134.v1.0.0).

## List of abbreviations

AAV: adeno-associated virus
ACB: nucleus accumbens
AD: Alzheimer’s disease
BLA: baso-lateral amygdala
BOLD: blood-oxygen-level-dependent
DG: dentate gyrus
DMN: default-mode network
ENTl: lateral entorhinal cortex
ILA: infralimbic area
mPFC: medial prefrontal cortex
ofMRI: optogenetics functional MRI
ReHo: regional homogeneity
rsfMRI: resting state functional MRI

## Authors’ contributions

FM: Conceptualization, Formal analysis, Investigation, Methodology, Validation, Visualization, Writing - original draft, Writing - review and editing, Funding acquisition.

LYY, RB, LS, HGB, SHB, CYL, HM, CLT, JHL: Validation, Writing - Review & Editing

RB: Software, Methodology, Writing - Review & editing.

CH: Writing - reviewing and editing, Analysis interpretation

FY, SY, MKPL: Validation, Resources, Writing - Review & Editing, Funding acquisition.

MO: Resources, Writing - Review & Editing, Supervision, Funding acquisition.

JGi: Conceptualization, Resources, Writing - Review & Editing, Supervision, Funding acquisition.

JGr: Conceptualization, Methodology, Formal analysis, Data curation, Software, Resources, Writing - Review & Editing, Supervision, Funding acquisition.

## Acknowledgements

This work was supported by the Singapore BioImaging Consortium core funding, Singapore BioImaging Consortium award #2017 to FM and #2016 to JGr. FM was supported by a Ph.D. scholarship funded through the A*STAR Research Attachment Programme and the University of Manchester (awarded to JGi). This work was also supported by the A*STAR Investigatorship (awarded to FY) and JCO grant (BMSI/15-800003-SBIC-OOE) awarded to SY.

