## Supplemental material for "The lateral entorhinal cortex is a hub for local and global dysfunction in pre-tauopathy states"

### Supplementary Material

#### Supplementary Methods

##### Electrophysiological recordings *in vivo*: anesthesia and surgical procedure

Anesthesia was induced via intraperitoneal (i.p) injection of urethane (1.5-1.7 g/kg of 30% w/v solution prepared in 0.9% saline; ethyl carbamate, Sigma, UK). An additional dose of urethane (50 µl of 10% w/v urethane solution prepared in 0.9% saline) was administered if complete areflexia was not reached after 40 minutes from the first injection. Mice were mounted and fixed in a stereotaxic frame and mouse adapter (Kopf 1430, USA) to immobilize the head prior to surgery. Body temperature was maintained around 37°C throughout the duration of the experiment with the use of a homoeothermic blanket (Harvard, UK) and a rectal thermistor probe placed underneath the abdomen. A midline scalp incision was made, and the skin retracted to expose the skull. After identifying Bregma and Lambda, the distance between these two landmarks was measured and compared to the standard 4.2 mm reported in the mouse brain atlas [57]. Any deviation from 4.2 mm allowed a proportional adjustment for craniotomy coordinates to improve the accuracy of the electrode placement. Craniotomies were made above the left hemisphere using a 0.9 mm drill bit (Fine Science Tools, Germany) and a high-speed handheld drill (Foredom, USA). Care was taken to maintain the craniotomy sites moist during the whole surgical procedure. Coordinates for craniotomy and electrode placement were marked onto the skull, relative to Bregma and the midline for recording in BLA (Bregma: AP -2.5 mm, ML: +2 mm) and DG (Bregma: AP -3.5 mm, ML: +2.5 mm), as well as stimulation of ENTl (Bregma: AP -2.8 mm, ML: +4.2 mm). Recording electrodes consisted of 32-contact probes, each laid out as 2 shanks of 16 electrodes that were 500 µm apart with 100 µm between recording points (A2x16-10-500-100-413, NeuroNexus Technologies, MI). These were first lowered to target locations (BLA target was 4 mm ventral from the brain surface at a 15° angle from vertical in the coronal plane; the DG target was also 4 mm ventral to brain surface but along a vertical plane) and then used to record brain activity during different stimulation protocols. The 2 recording electrode shanks were coated in Vibrant CM-DiI (Sigma, UK) cell-labeling solution to allow post-mortem localization of their location with fluorescence microscopy. The stimulation electrode consisted of twisted, diameter 125 µm Teflon-insulated stainless steel wires (Advent RM, UK) and was inserted an initial 3 mm from the brain surface in one step and then slowly lowered further during continuous application of stimulating pulses (50 ms PPS, 300 μA, 0.2 ms duration pulses) until the expected responses in DG and BLA were seen on an oscilloscope.

**Electrophysiological recordings *in vivo*: data acquisition**

Data were recorded on a Recorder64 system (Plexon Inc, USA), and saved for offline analysis. Electrodes were connected to a head stage (fixed gain of x20) and then to a preamplifier for a total gain of x500. Field excitatory postsynaptic potentials (fEPSPs) were recorded at a sampling rate of 5 or 10 kHz using a 12-bit A/D converter and then stored for offline analysis. A low pass filter (1 kHz) was applied to attenuate spiking activity. Electrical stimuli were delivered by a constant-current stimulator (DS3, Digitimer, UK), triggered by analog 5 V square wave pulses from a National Instruments PCI card (PCI-6071E). Timings and types of stimuli to be delivered were controlled through custom-written programs in LabVIEW (v8, National Instruments). Stimulus duration was fixed at 200 μs throughout each protocol. Two different protocols of stimulation were performed: Input/Output, for the Input/Output curve (IOC) analysis and PPS for the PPI analysis. In each mouse, the channel with the most distinctive response (as revealed through current source density analysis *post hoc*; see below) was selected for further analysis.

**Electrophysiological recordings *in vivo*: stimulation protocols - Input/output curve**

The IOC reflects the functional strength of synaptic connectivity: by applying different current intensities, it is possible to analyze how the response (Output voltage) changes as a function of input strength (Input current). The range of current intensities used here was 180, 300, 450 and 600 μA.

**Electrophysiological recordings *in vivo*: stimulation protocols - Paired-pulse stimulation**

To measure short-term synaptic plasticity, stimulus current was set to half-maximum of the response obtained in the BLA IOC paradigm, i.e. ~300 μA, and paired-pulses were delivered at this current with different paired-pulse intervals for 20 repetitions each. The range of intervals was 20, 50, 100, 200, 500 and 1000 ms.

**Electrophysiological recordings *in vivo*: data analysis**

For the IOC protocol, stimulation at each current intensity was repeated 20 times (runs), the initial slope of the fEPSP response measured for each repetition and then these 20 values were averaged. The mean response to each current step for each mouse was used to plot the IOC of response to current intensity by genotype, age, and ROI. For the PPS protocol, the slope for the initial fEPSP response on a selected channel was measured for both stimuli in each pair. For each pair, the fEPSP response to the second stimulus (P2) was normalized to that of the first (P1) and expressed as PPI ratio (see equation 1, below), such that positive values indicate synaptic paired-pulse facilitation (PPF) and negative values synaptic PP depression (PPD).

$$PPI = (P2 - P1) / P1 \qquad (1)$$

In the case of synaptic facilitation PPI > 0, whereas, PPI < 0 for synaptic depression. For both the IOC and PPS results, statistical analysis was performed using the statistical software R 3.6.1 (The R Foundation for Statistical Computing). Responses to the IOC (initial slope for P1) and PPI (ratio between P2 and P1) paradigms in BLA and DG at the two age points were analyzed separately with a linear mixed model analysis, using ROI, age, genotype, amplitude/intervals (amplitude for IOC and interval for PPS), and their mutual interactions as fixed effects and animal intercepts and run repeats as random effects using R package lme4. Interaction effects were determined using a likelihood ratio test. A *post hoc* analysis was performed after a general linear hypothesis test using contrast to determine genotype effects at each stimulation amplitude or interval. Correction for multiple comparisons was implemented during the contrast analysis using the R package multcomp [69,70].

Data are plotted as mean ± 1 standard deviation (SD).

**Brain slice preparation and whole-cell current-clamp recordings in ILA and DG**

After undergoing the scan at 6 months, 8 mice ($N_{controls}$ = 4, $N_{3xTgAD}$ = 4) from the ofMRI experimental mice were used for a whole-cell patch recording in brain slices. The animals were deeply anesthetized with ketamine/xylazine (0.1 ml/kg) and cardiac perfusion was performed with ice-cold, oxygenated

(95% $O_2$ and 5% $CO_2$) NMDG-HEPES solution consisting of 93 NMDG, 2.5 KCl, 1.2 $NaH_2PO_4$, 30 $NaHCO_3$, 20 HEPES, 25 glucose, 5 sodium ascorbate, 2 thiourea, 3 sodium pyruvate, 10 $MgSO_4$, 0.5 $CaCl_2$ (in mM, pH 7.3-7.4, 300-310 mOsm). After perfusion, the brain was coronally sliced at 350 µm thickness using a VT-1200S vibratome (Leica, Germany) in ice-cold, oxygenated NMDG-HEPES solution. Brain slices were transferred to the pre-warmed NMDG-HEPES solution and left to recover for 35 min with constant oxygenation at 37℃. During this recovery, 250, 250, 500, 1000, and 2000 µl of 2 M NaCl solution were added at 10, 15, 20, 25, 30 min recovery time points, respectively, into 150 ml of the recovery solution. After recovery, brain slices were placed into HEPES-holding solution consisting 92 NaCl, 2.5 KCl, 1.2 $NaH_2PO_4$, 30 $NaHCO_3$, 20 HEPES, 25 glucose, 5 sodium ascorbate, 2 thiourea, 3 sodium pyruvate, 2 $MgSO_4$, 2 $CaCl_2$ (in mM, pH 7.3-7.4, 300-310 mOsm)[71] at room temperature.

For recording, brain slices were transferred to a recording chamber perfused with CSF solution consisting of 124 NaCl, 2.5 KCl, 1.2 $NaH_2PO_4$, 24 $NaHCO_3$, 5 HEPES, 12.5 glucose, 2 $MgSO_4$, 2 $CaCl_2$ (in mM, pH 7.3-7.4, 300-310 mOsm) at room temperature. Recording pipettes were prepared from borosilicate glass pipette using a P-1000 puller (Sutter Instrument, USA) to 4-7 MΩ resistance. Current-clamp recording was then performed in the infralimbic cortex layer 2/3 and hippocampal dentate gyrus using recording pipette filled with an internal solution consisting 130 K-gluconate, 0.1 EGTA, 1 $MgCl_2$, 10 HEPES, 5 NaCl, 11 KCl, 5 phosphocreatine, 2 Mg-ATP, 0.3 Na-GTP (in mM, pH 7.3-7.4, 300-310 mOsm). Data were collected and recorded by Multiclamp 700A amplifier (Axon Instruments, USA), Digidata 1550B (Axon Instruments, USA), pCLAMP v10 (Molecular Devices, USA), and HEKA ECP10 USB (HEKA Elektronik, Germany), PatchMaster v2x90.2. Only data that met our criteria (leakage current, <100 pA; R-series, <30 MΩ) were used for further analysis. Neuronal intrinsic properties were analyzed using AxoGraph X with statistical analysis conducted using Prism 7.0 (GraphPad).

**Whole-cell patch recording of acute brain slices**

Mouse brains were rapidly removed after decapitation and placed in high sucrose ice-cold oxygenated artificial cerebrospinal fluid (ACSF) containing the following (in mM): 230 sucrose, 2.5 KCl, 10

$MgSO_4$, 0.5 $CaCl_2$, 26 $NaHCO_3$, 11 glucose, 1 kynurenic acid, pH 7.3, 95% $O_2$ and 5% $CO_2$. Coronal brain slices were cut at a thickness of 250 μm using a vibratome (VT1200S; Leica Biosystems) and immediately transferred to an incubation chamber filled with ACSF containing the following (in mM): 119 NaCl, 2.5 KCl, 1.3 $MgCl_2$, 2.5 $CaCl_2$, 1.2 $NaH_2PO_4$, 26 $NaHCO_3$, and 11 glucose, pH 7.3, equilibrated with 95% $O_2$ and 5% $CO_2$. Slices were allowed to recover at 32°C for 30 minutes and then maintained at room temperature. Experiments were performed at room temperature. Whole-cell patch-clamp recordings were performed on ENTl pyramidal cells expressing ChR2-mCherry and were visualized using a CCD camera and monitor. Pipettes used for recording were pulled from thin-walled borosilicate glass capillary tubes (length 75 mm, outer diameter 1.5 mm, inner diameter 1.1 mm, WPI) using a DMZ Ziets-Puller (Zeitz). Patch pipettes (2–4 MΩ) were filled with internal solution containing (in mM): 105 K-gluconate, 30 KCl, 4 $MgCl_2$, 10 HEPES, 0.3 EGTA, 4 Na-ATP, 0.3 Na-GTP, and 10 $Na_2$-phosphocreatine (pH 7.3 with KOH; 295 mOsm), for both voltage- and current-clamp recordings. Photostimulation (460 nm) was delivered by an LED illumination system (pE-4000). Several trains of a square pulse of 20 ms duration with 5, 10, and 20 Hz, were delivered respectively under current-clamp mode (I = 0) to examine whether the neurons were able to follow high-frequency photostimulation. After different frequencies of photostimulation were completed, neurons were shifted to voltage-clamp mode (at -60 mV), and a prolonged square pulse of 500 ms duration was delivered, to further confirm whether ChR2-induced current could be seen in the recorded neurons. The access resistance, membrane resistance, and membrane capacitance were consistently monitored during the experiment to ensure the stability and the health of the cell.

**ELISA diagnostic assay**

Brains were homogenized in a Tris-HCl buffer and agitated for 30 minutes before being subjected to centrifugation at 6000 g. The supernatant was used for the detection of soluble $A\beta_{40\text{-}42}$ and the pellet was re-suspended in Tris-HCl and 10 μl was used for further processing for insoluble Aβ ELISA as recommended by the ELISA kit (incubation with 5 M Guanidine to solubilize any aggregates and diluted before adding into ELISA plates). Samples were diluted only when necessary (phospho-tau and total tau were diluted 2x, no dilution for Aβ ELISAs). As readings could be affected by the sample volume in each well, we normalized the ELISA results to protein assay results of the same fraction, hence, the final units were in picogram of Aβ or tau in per milligram of protein (pg/mg protein).

##### Immunohistochemistry for AT8 and 6e10

Immunohistochemical analyses were performed as described previously in [72]. Brains of $N_{controls}$ = 6 (2 per age point: 3, 6 and 10 months of age) and $N_{3xTgAD}$ = 6 (2 per age point: 3, 6 and 10 months of age) were fixed by perfusion with 4% paraformaldehyde (PFA) in 0.01 M phosphate buffer saline (PBS) and then in PFA and 30% sucrose for 48 h at 4°C. Fixed brains were cut on a microtome (CM3050S, Leica Microsystems, Nussloch, Germany) into 45 µm-thick sections and collected into a cold cryoprotectant solution (80 mM $K_2HPO_4$, 20 mM $KH_2PO_4$, 154 mM NaCl, 0.3 g/ml sucrose, 0.01 g/ml polyvinylpyrrolidone, 30% vol/vol ethylene glycol). Sections were washed 5 times for 3 minutes in 1 x PBS and blocked in 5% FBS with 0.1% Triton X-100 for 1 hour, followed by overnight incubation with the primary antibody in blocking solution at 4°C. These brain sections were then immunostained with primary antibodies against Aβ (6e10, Covance Research Products Inc Cat #SIG-39300-1000 RRID: AB_10175637) and phospho-tau (AT8 ThermoFisher Scientific, Cat. #MN1020). After the primary antibody binding step sections were washed 5 times in 1 x PBS for 3 min and then incubated with anti-mouse Alexa 488 or anti-rabbit Alexa 594 for 2 hours followed by washing 3 times with 1 x PBS for 3 minutes. Sections were then mounted with DAPI plus mounting media on slides. All pictures were taken by using a confocal microscope with a 40× objective.

##### *Ex vivo* histology

After the completion of the experiments, animals were injected with an overdose of ketamine/xylazine and transcardially perfused with PBS (0.01 M) followed by 4% PFA in 0.01 M PBS. After extraction, the brain was post-fixed in 4% PFA overnight. Brain sections of 50 µm were made with a vibratome (VBT1200s, Leica) and fluorophore expression, together with Hoechst staining, was checked through a confocal microscope Ti-E; DS-Qi2; Fluorescence, SBIC-Nikon Imaging Center, Singapore) for anatomical confirmation of viral injection and fiber optic cannula positioning (Figure 2).

#### Supplementary Figures

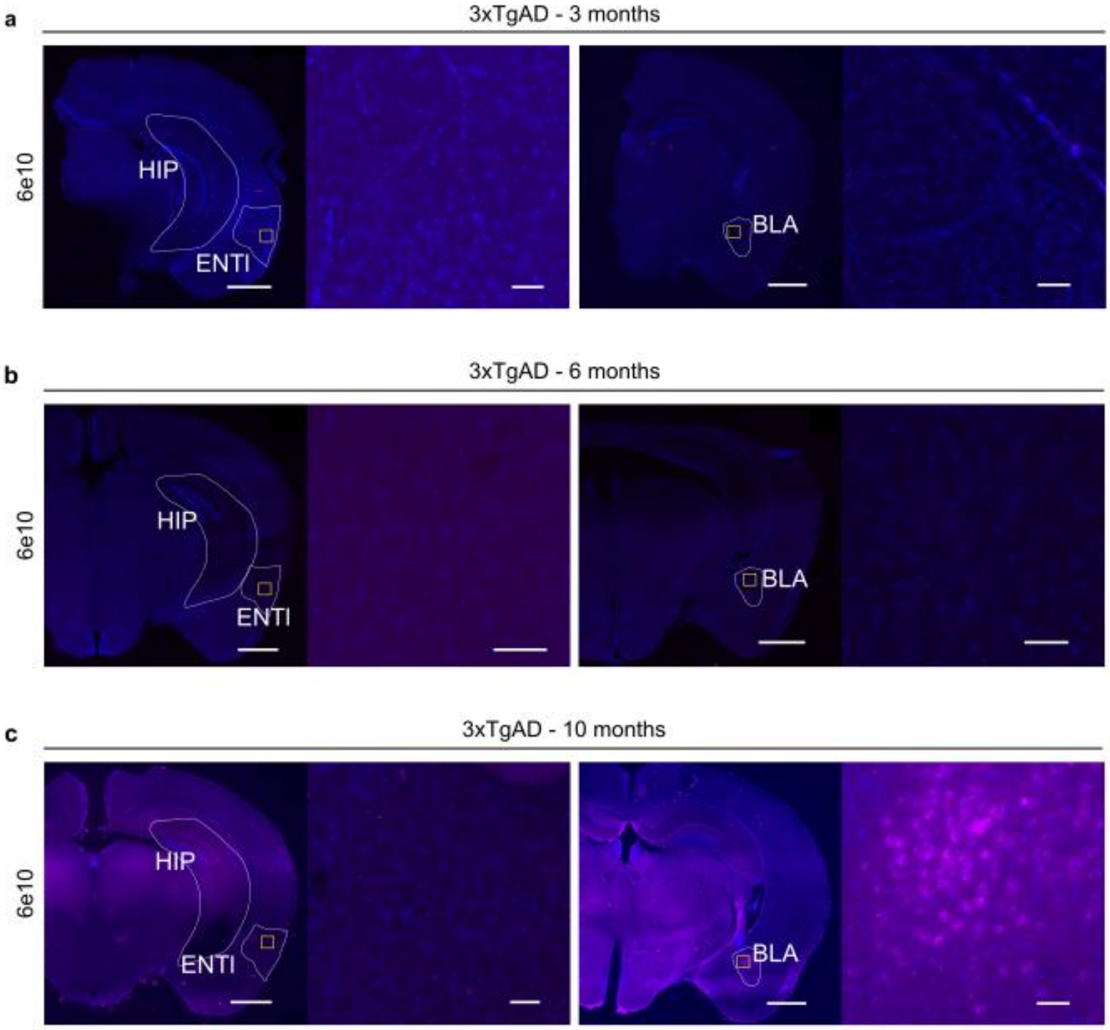


**Figure S1: Immunohistochemistry for 3xTgAD characterization with 6e10-DAPI.**

3xTgAD mice show no reactivity to the N-terminus of Aβ form at both 3 and 6 months of age in the hippocampus (HIP), ENTl (left panels, lower and higher magnification, left to right) and BLA (right panels, lower and higher magnification, left to right) **a,b)**. **c)** 3xTgAD mice show Aβ-positive labeling in the BLA at 10 months, right panels, lower and higher magnification, left to right. Lower magnification: 1000 µm; higher magnification: 100 µm.

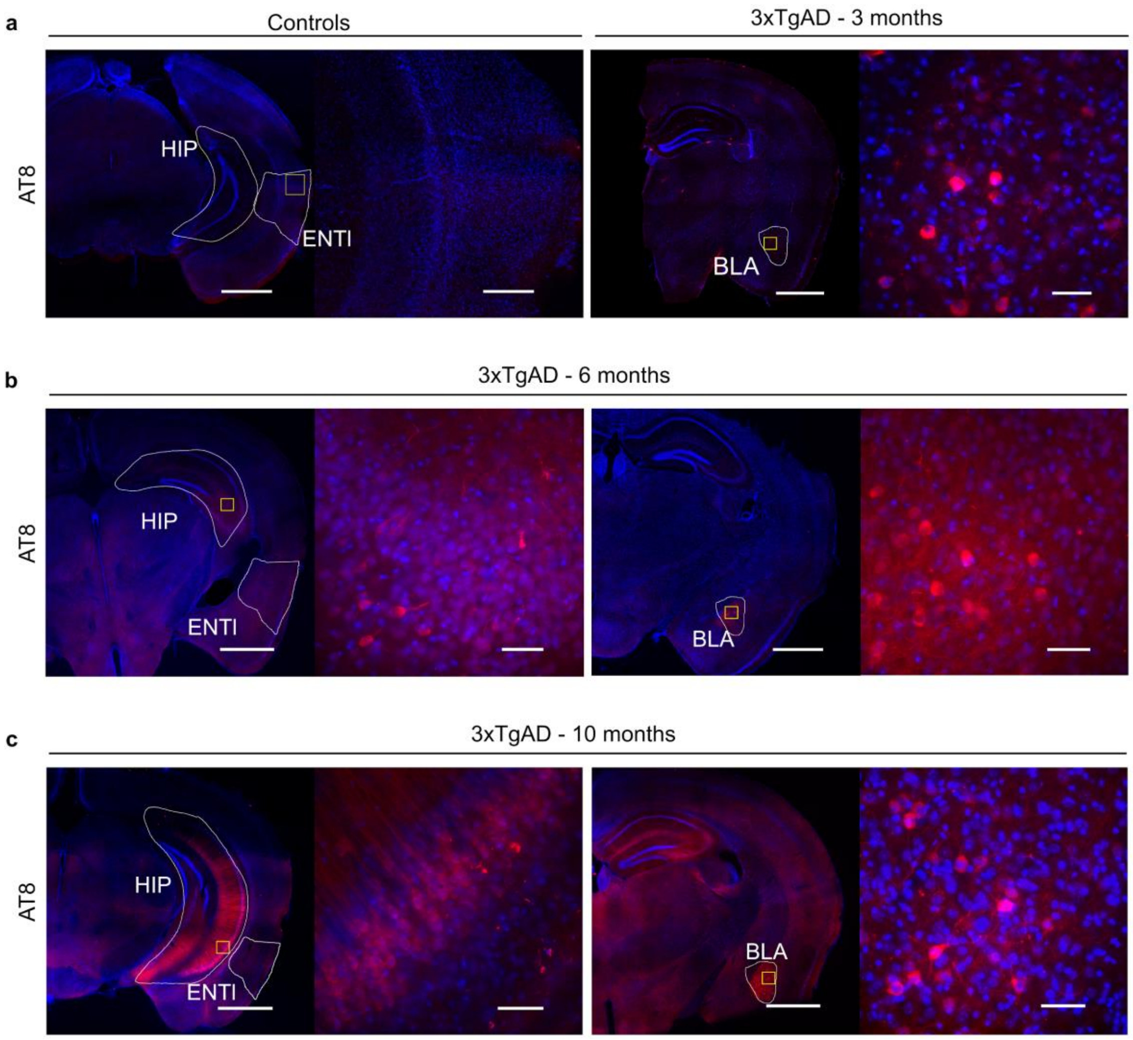


**Figure S2: Immunohistochemistry for 3xTgAD characterization with co-labeling AT8-DAPI**

**a)** No reactivity was detected in wild-type mice (3-month-old example, left panels with lower and higher magnification, left to right), whereas 3xTgAD mice showed positive staining for phosphorylated tau in the amygdala area by 3 months of age (right panels with lower and higher magnification, left to right). **b)** 6-month-old 3xTgAD mice show tau-positive labeling extended to the hippocampus (HIP, left panels, lower and higher magnification, left to right), whereas ENTl shows no apparent pathology; tau-positive labeling is confirmed in the amygdala (BLA, right panels, lower and higher magnification, left to right). **c)** 10-month-old 3xTgAD mice show tau-positive staining in the hippocampus (left panels, lower and higher magnification, left to right). ENTl, however, does not show a positive response to AT8. Confirmed tau-positive labeling was seen in the amygdala (right panels, lower and higher magnification, left to right). Lower magnification: 1000 µm, higher magnification: 100 µm

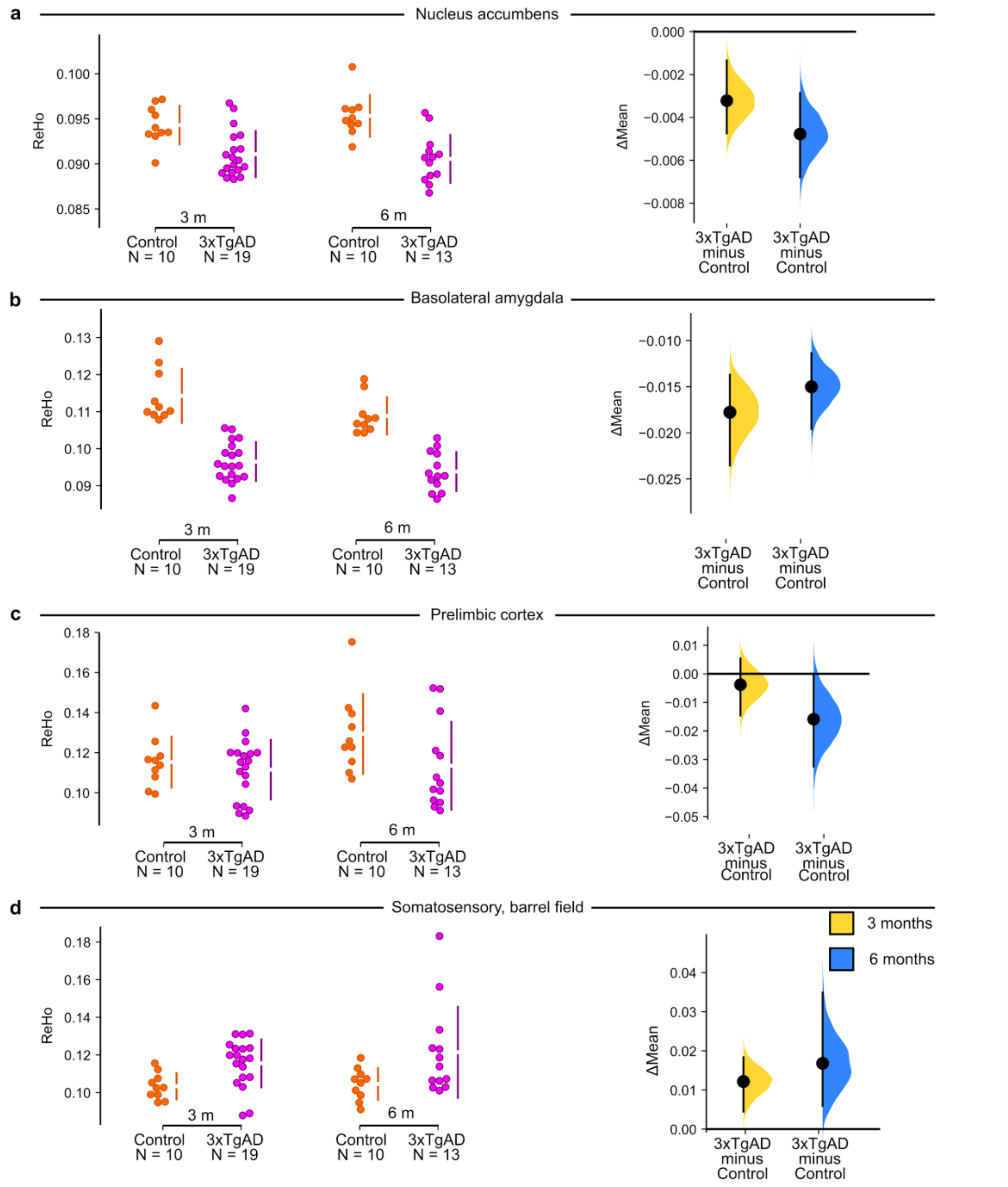


**Figure S3 ReHo values extracted in selected ROIs show functional connectivity changes in 3xTgAD**

3xTgAD mice show deficits in ReHo in regions encompassing **a)** the ACB ($\Delta mean_{3months}$ = -0.110 [-0.154, 0.068], $\Delta mean_{6months}$ = -0.152 [-0.199, -0.107]); **b)** the BLA ($\Delta mean_{3months}$ = -0.066 [-0.110, -0.025], $\Delta mean_{6months}$ = -0.061 [-0.115,-0.011]); **c)** the mPFC, prelimbic reported as example ($\Delta mean_{3months}$ = -0.004 [-0.015, -0.005], $\Delta mean_{6months}$ = -0.016 [-0.032, 1.93E-05]); **d)** increase in ReHo was found in somatosensory areas (SSp-bfd, $\Delta mean_{3months}$ = 0.0833 [0.0192, 0.149], $\Delta mean_{6months}$ = 0.112 [0.0587,0.157]) suggests that functional connectivity deficits in 3xTgAD might be network-specific to AD-vulnerable regions. ACB: nucleus accumbens; BLA: basolateral amygdala; ReHo: regional homogeneity

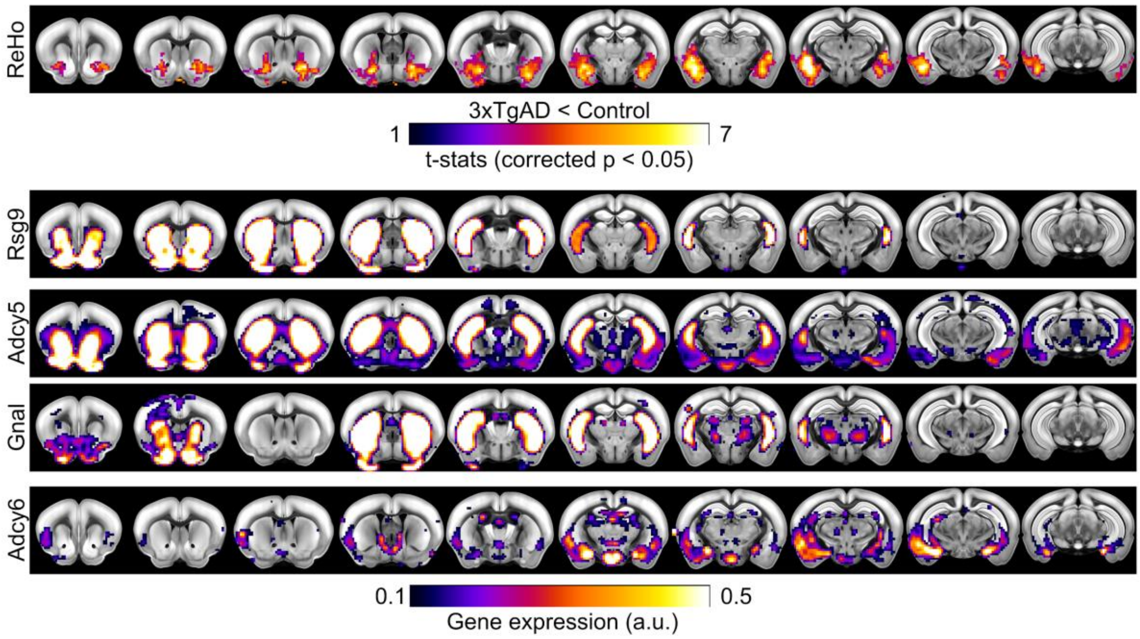


**Figure S4: Correlated gene expression.**

Selected gene expression maps derived from the Anatomic Gene Expression Atlas present mild correlations (r range [0.2 - 0.14]) with ReHo deficits in 3 months old 3xTgAD.

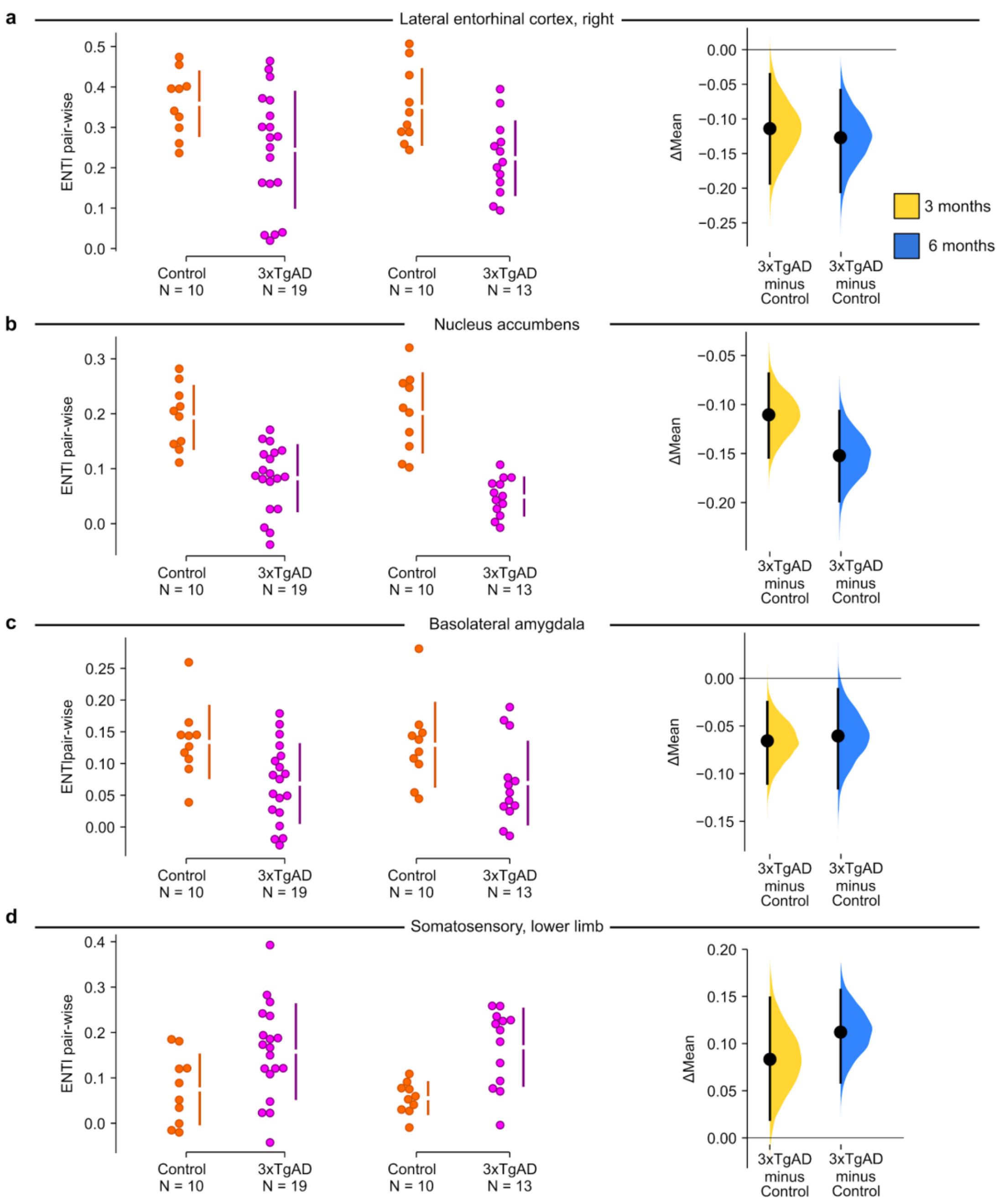


**Figure S5: Pair-wise ROI interactions relative to the left-hemisphere ENTl.**

Compared to the left ENTl, 3xTgAD mice show a decrease in functional connectivity in the right ENTl **(a)**, ACB **(b)** and BLA **(c)**. These results show consistent trends as in whole-brain ReHo, highlighting the relevance of ENTl as a central hub for functional connectivity changes in 3xTgAD. **d)** Functional connectivity in somatosensory regions is increased in 3xTgAD compared to controls, in pair-wise interaction with left ENTl.

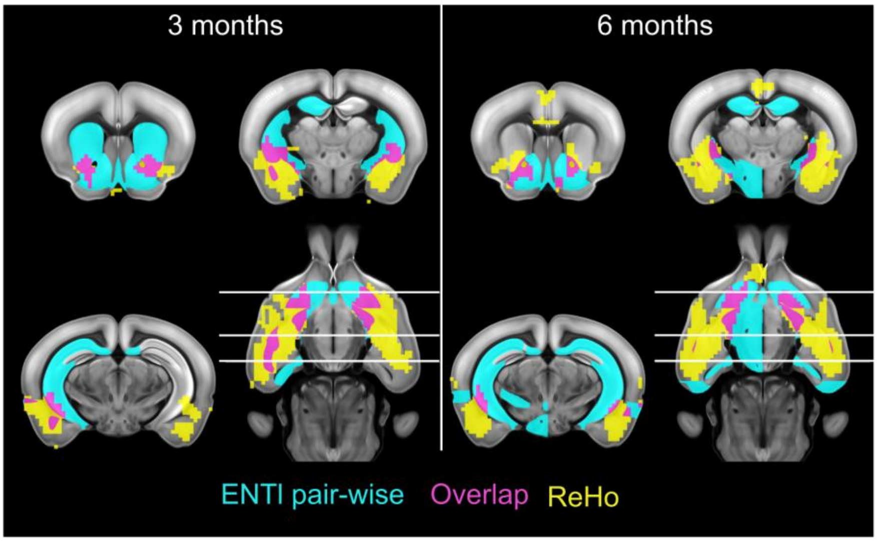


**Figure S6: Restricted pair-wise ENTl at 3 and 6 months overlapped with ReHo.**

The overlap (pink) between ReHo (yellow) and pair-wise interactions between ENTl and whole-brain regions (cyan) resulted in major hotspots of interest, consistent across ages.

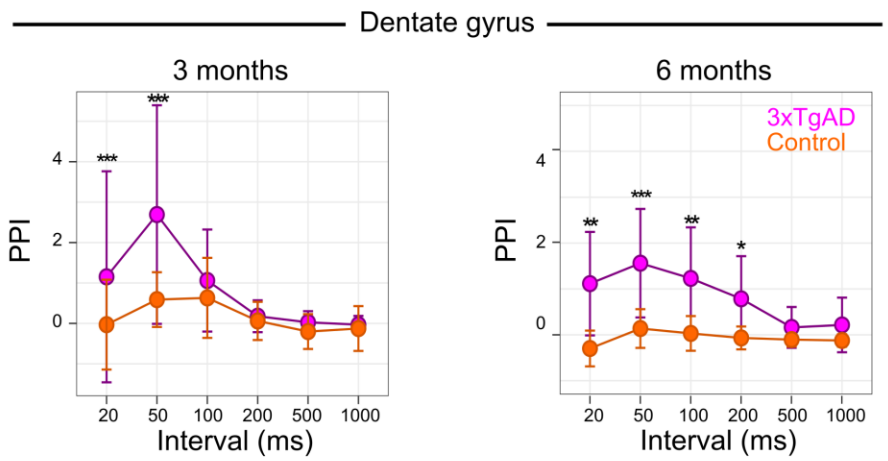


**Figure S7: Paired-pulse stimulation in DG at 3 and 6 months shows increased facilitation in 3xTgAD.**

Response to ENTl stimulus pairs (PPS) at paired-pulse intervals of 20, 50, 100, 200, 500, and 1000 ms. Field EPSPs response to PPS recorded in DG in 3 (left) and 6 (right) months old 3xTgAD mice and matched controls showed a significant increase in facilitation at short intervals: 20 ms, 50 ms. DG shows a genotype effect at intervals of 100 and 200 ms. Data are plotted as mean ± SD. * $p < 0.05$, ** $p < 0.005$, *** $p < 0.001$. PPS: paired-pulse stimulation. ENTl: lateral entorhinal cortex, BLA: basolateral amygdala; DG: dentate gyrus; fEPSP: field excitatory postsynaptic potential.

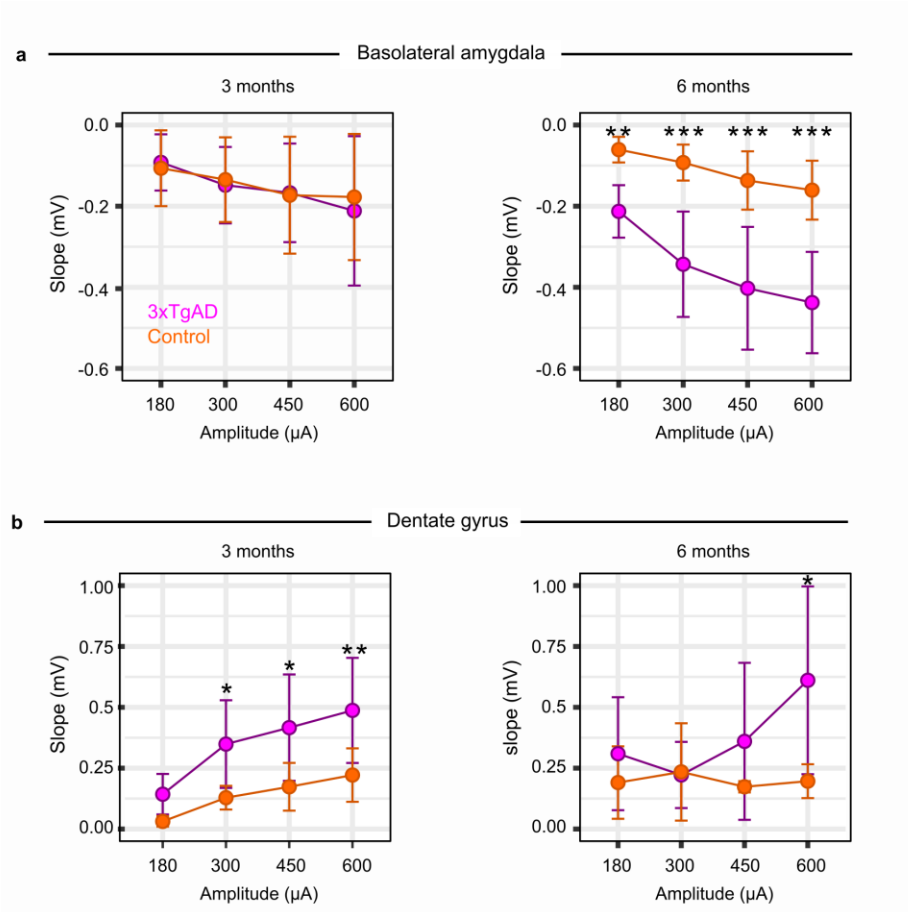


**Figure S8: Input/Output curves for BLA and DG response to ENTl stimulation at 3- and 6-months show strengthening of synaptic connectivity in 3xTgAD.**

**a, b)** Input/output curve for 3 (left) and 6 (right) month old 3xTgAD mice and matched controls in BLA and DG, respectively. Data are plotted as mean initial fEPSP slope values ± SD. * $p < 0.05$, ** $p < 0.005$, *** $p < 0.001$. ENTl: lateral entorhinal cortex, BLA: basolateral amygdala; DG: dentate gyrus.

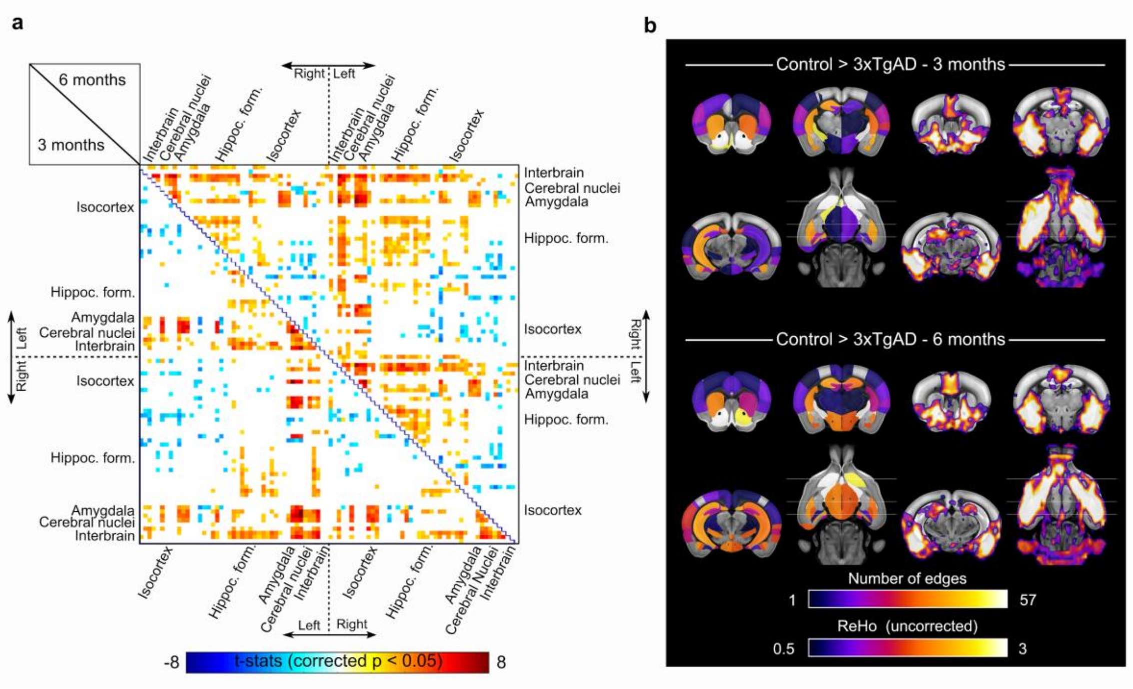


.

**Figure S9: Whole-brain NA at 3 months and 6 months and overlap with ReHo analysis.**

**a)** Half-matrices of whole-brain network analysis show deficits in 3xTgAD at 3 and 6 months of age. **b)** Overlap of ReHo results (uncorrected) with NA spatial maps highlight common regions showing deficits in functional connectivity, such as the amygdaloid/cerebral nuclei (e.g. BLA and ACB) and hippocampal formation (e.g. ENTl). ReHo: regional homogeneity; NA: network analysis; ENTl: lateral entorhinal cortex; BLA: basolateral amygdala; ACB: nucleus accumbens.

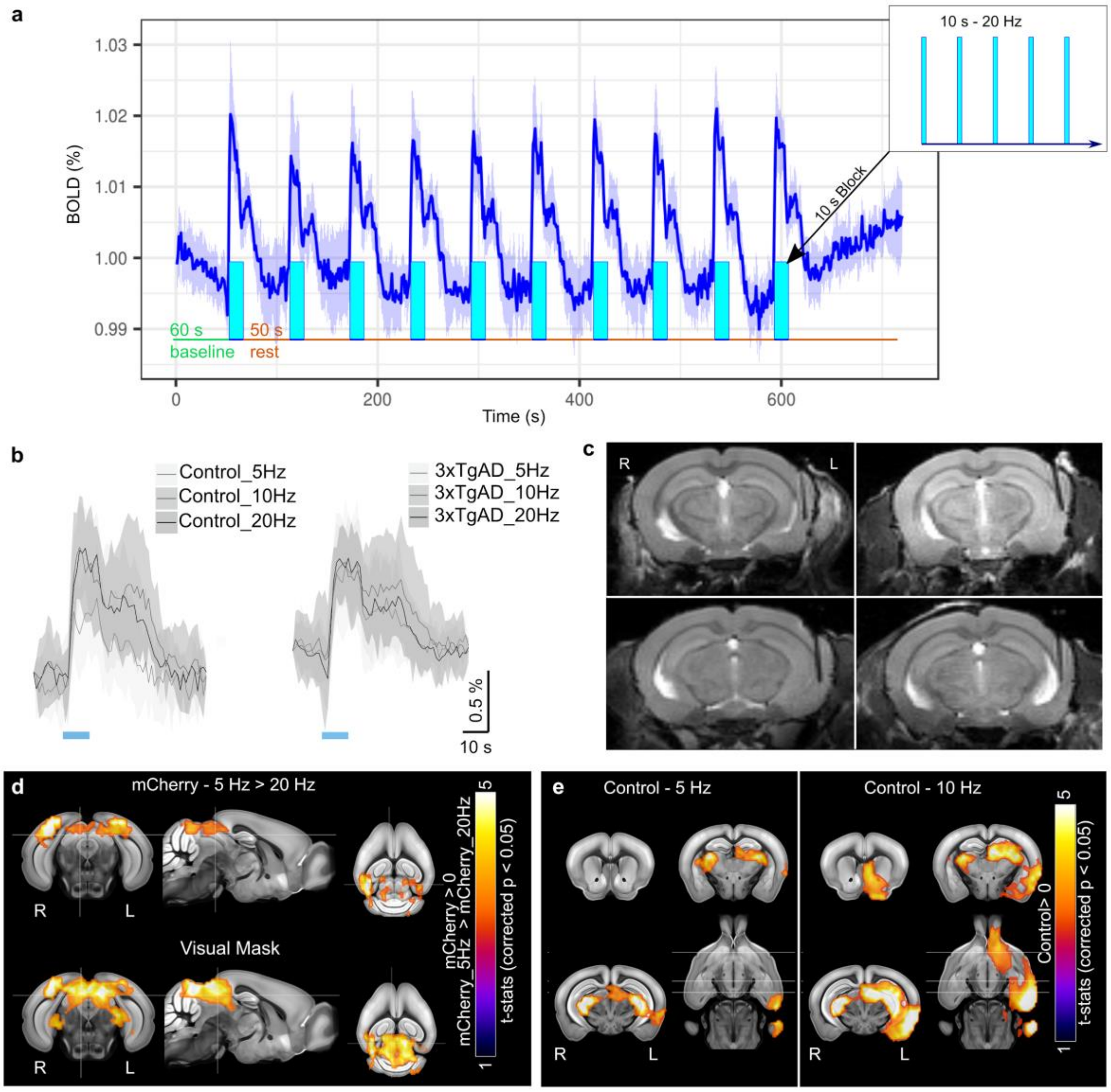


**Figure S10: Experimental design for optogenetic stimulation and control validation.**

**a)** Stimulation design consisted of 50 s baseline and subsequent 10 blocks of 5, 10, or 20 Hz-pulses for 10 s, alternated by 50 s of rest with no stimulation; stimulation protocols were provided in random order. **b)** Average BOLD response for the 10 blocks of stimulation across frequencies for controls and 3xTgAD (left and right respectively), represented as mean ± SD. **c)** The anatomical scan shows the fiber positioning in 4 mice from the experimental group. **d)** Top: Two-sample t-test in mice transfected with mCherry shows that only lower frequencies, i.e. 5 Hz, engage a stronger response in the visual system, compared to 20 Hz stimulation. Bottom: Response derived from a 5 Hz stimulation protocol was used to mask the regions that responded positively to visual stimulation. **e)** Mice injected with ChR2 show response to low frequencies, i.e. 5 and 10 Hz, here shown as the control group at 3 months of age. The response to 20 Hz stimulation was chosen for the rest of the analyses.

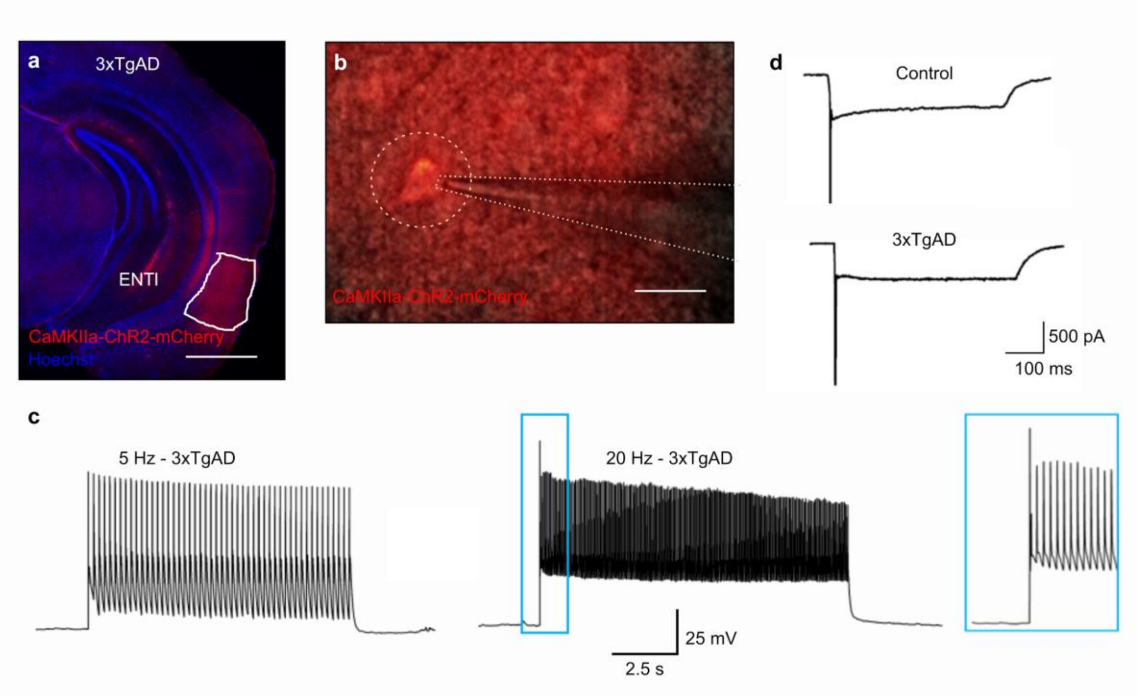


**Figure S11: Optogenetic validation of pyramidal neurons in ENTl.**

**a)** Histological evidence for opsin expression in 3xTgAD mice co-stained with Hoechst neuronal cell bodies. Scale bar: 1000 µm.**b)** whole-cell patch-clamping in a ChR2-expressing pyramidal neuron in the ENTl. Scale bar: 25 µm. **c)** 5 and 20 Hz stimulation of the patched neuron shows faithful frequency-gated activity. **d)** Prolonged photostimulation (500 ms) applied under voltage-clamp mode (-60 mV) shows an initial action potential spike and a ChR2-induced inward current.

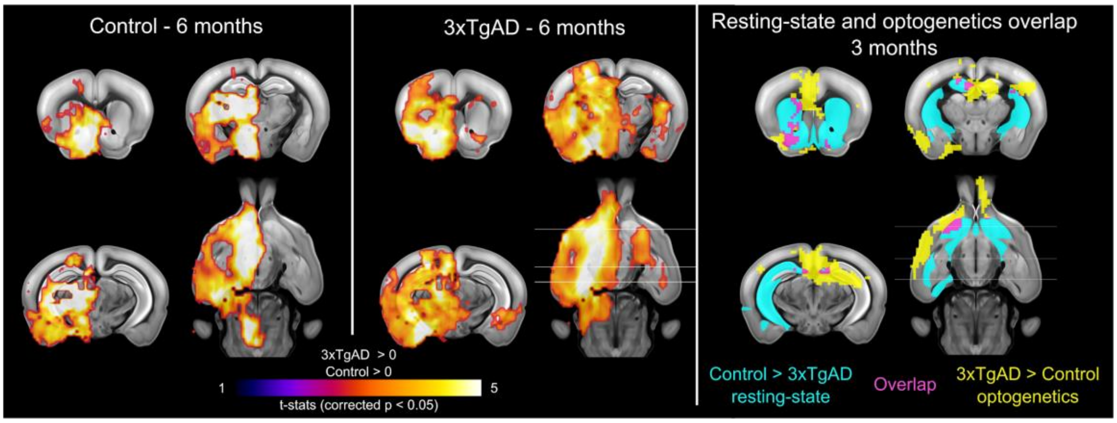


**Figure S12: Optogenetically-locked BOLD response in control and 3xTgAD mice at 6 months and the overlap with resting-state functional connectivity at 3 months.**

Left and middle panels indicate the one-sample t-test for stimulation-locked BOLD response in the control and 3xTgAD cohorts, respectively ($p < 0.05$ corrected), highlighting activation in key regions related to ENTl projections, i.e., HIP, BLA, ACB. The right panel indicates resting-state functional connectivity analysis (cyan) and optogenetically-evoked response (yellow) highlights overlapping *loci* for the decrease in functional connectivity at rest and potentiated response during stimulation (pink) at the 3-month-old age point.

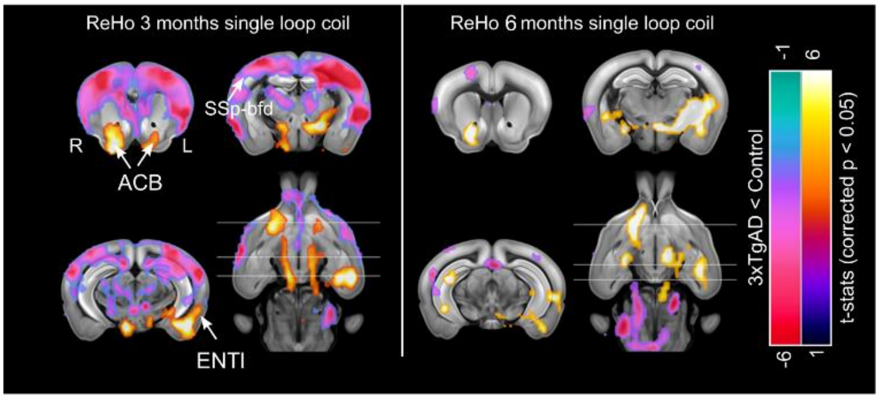


**Figure S13: Resting-state fMRI with single-loop coil reveals similar patterns of functional connectivity deficits in 3xTgAD.**

ReHo analysis of rsfMRI with the single-loop coil, acquired before the ofMRI protocols, reveals similar loss of coherence in AD-vulnerable brain regions, e.g., ENTl, ACB. The increase in ReHo in somatosensory areas is also in line with the results reported with the cryogenic coil. 3 months: $N_{controls} = 10$; $N_{3xTgAD} = 12$; 6 months: $N_{controls} = 8$; $N_{3xTgAD} = 10$. ENTl: lateral entorhinal cortex; ACB: nucleus accumbens; SSp-bfd: somatosensory barrel field cortex; ReHo: regional homogeneity.

**Table S1: Animal breakdown for the experimental procedures.**

| | 3 months | | 6 months | | 10 months |
|---|---|---|---|---|---|
| | Control | 3xTgAD | Control | 3xTgAD | 3xTgAD |
| RsfMRI | 10‣ | 19‣‣ | 10‣ | 13‣‣ | - |
| OfMRI-ChR2 | 10• | 12•• | 8• | 10•• | - |
| OfMRI-mCherry | 9 | - | - | - | - |
| PPS | 7 | 6 | 4 | 4 | - |
| IOC | 5 | 4 | 4 | 3 | - |
| Slice electrophysiology | | | 4• | 5•• | - |
| Slice optogenetics | 1 | 1 | - | - | - |
| Viral expression histology | 1 (ChR2-mCherry) •<br>1 (mCherry) | 1•• | - | - | - |
| Immunohistochemistry | 2 | 2 | 2 | 2 | 2 |
| ELISA | 2• | 1•• | 2• | 4•• | 1 |

RsfMRI and ofMRI experiments were conducted on different cohorts of animals. Within each cohort, longitudinal experiments between 3- and 6-months age points were performed. Subgroups of the experimental animals that previously underwent fMRI experiments, were then used for slice electrophysiology, viral expression validation, and ELISA tests. The same symbol represents animals belonging to the same cohort or a subgroup of the same cohort. Electrophysiological experiments *in vivo* were conducted on different cohorts, being this a non-recovery procedure. Immunohistochemistry stainings were performed on a separate sample of mice.

**Table S2: Intrinsic properties of granule cells in the dentate gyrus and pyramidal cells in the infralimbic cortex, part of the medial prefrontal cortex.**

| | **Dentate gyrus** | | | **Infralimbic cortex** | | |
|---|---|---|---|---|---|---|
| | Controls N = 4, n = 29 | 3xTgAD N = 5, n = 18 | p-value (M-W) | Controls N = 4, n = 10 | 3xTgAD N = 5, n = 11 | p-value (M-W) |
| Rm (mV) | -74.54±0.7242 | -76.66±1.029 | ns | -52.42±2.827 | -54.71±3.149 | ns |
| Rheobase (pA) | 79.83±6.231 | 69.44±6.591 | ns | 37±8.95 | 24.55±3.659 | ns |
| Threshold (mV) | -33.38±1.554 | -36.38±1.919 | ns | -35.69±0.5727 | -34.58±1.228 | ns |
| AP latency (ms) | 238±40.01 | 343.3±56.88 | ns | 94.81±28.06 | 174.2±33.35 | ns |
| AP amplitude (mV) | 97.06±2.129 | 91.59±2.219 | ns | 72.99±2.284 | 70.34±2.871 | ns |
| AHP (mV) | -11.35±0.8297 | -10.17±1.241 | ns | -10.9±1.653 | -10.73±0.9142 | ns |
| AHP latency (ms) | 5.015±0.2014 | 4.794±0.3514 | ns | 16.88±3.091 | 55.6±13.21 | 0.0028 (**) |
| Half-width (ms) | 1.321±0.0424 | 1.242±0.0472 | ns | 2.27±0.1154 | 2.862±0.2157 | 0.0079 (**) |

Rheobase, threshold, action potential (AP) latency, AP amplitude, afterhyperpolarization (AHP), AHP latency, and AP half-width were measured from the first trace showing action potentials (Rheobase trace). N = number of animals, n = number of cells. The values represent mean value with SEM and the statistical significance by asterisks (* $p < 0.05$, ** $p < 0.01$ by Mann-Whitney U test).

**Table S3: Number of action potential evoked by various current injections.**

| | Dentate gyrus | | | Infralimbic cortex | | |
|---|---|---|---|---|---|---|
| Current injection (pA) | Controls N = 4, n = 29 | 3xTgAD N = 5, n=18 | p-value (M-W) | Controls N = 4, n = 10 | 3xTgAD N = 5, n = 11 | p-value (M-W) |
| 20 | na | na | na | 1.8±0.7572 | 2.182±0.6851 | ns |
| 40 | 0.1034±0.076 | 1.556±0.879 | ns | 4±1.229 | 6±0.9723 | ns |
| 60 | 0.5517±0.236 | 3.556±1.322 | 0.029 (*) | 6.2±1.724 | 9.636±0.8008 | ns |
| 80 | 2.621±0.5985 | 6.056±1.652 | ns | 7.8±1.825 | 11.45±0.9757 | ns |
| 100 | 4.345±0.7773 | 8.5±1.801 | ns | 9.2±1.611 | 12.55±1.012 | ns |
| 120 | 5.241±1.013 | 10.39±1.91 | 0.0214 (*) | 8±1.366 | 13.82±1.094 | 0.0036 (**) |
| 140 | 6.138±1.24 | 10.83±1.526 | 0.0078 (**) | 8.4±1.392 | 13.45±1.268 | 0.0204 (*) |

The number of action potentials was counted in granule cells in the dentate gyrus and pyramidal cells in the infralimbic cortex from the various current injections (20 – 140 pA). N = number of animals, n = number of cells. The values represent mean value with SEM and the statistical significance was presented with asterisks (* p <0.05, ** p <0.01 by Mann-Whitney U test).
